# Multi-Omics Reprogramming Drives a Counterintuitive Reversal of Disease Susceptibility During Ageing

**DOI:** 10.64898/2025.12.12.693928

**Authors:** Alejandro Valdivieso, Léo Duperret, Bruno Petton, Gaelle Courtay, Océane Romatif, Juliette Pouzadoux, Sylvain Henry, Andrei Turtoi, Eve Toulza, Arnaud Lagorce, Lionel Degremont, Benjamin Morga, Emmanuel Vignal, Celine Cosseau, Fabrice Pernet, Guillaume Mitta, Jeremie Vidal-Dupiol

## Abstract

Ageing is a progressive and irreversible biological process characterized by the deterioration of physiological functions and increased vulnerability to mortality. Although extensively studied in vertebrates, ageing in long-lived invertebrates remains comparatively unexplored. While ageing typically leads to greater susceptibility to infectious diseases, a striking and unexpected reversal was identified in oysters: older oysters exhibit markedly increased tolerance to the Pacific Oyster Mortality Syndrome (POMS), a panzootic disease primarily driven by the OsHV-1 herpesvirus and responsible for severe losses in global aquaculture. To investigate this counterintuitive pattern, we challenged oysters aged 4, 16, and 28 months from four biparental families and conducted an integrative multi-omics analysis, including epigenomics, transcriptomics, and metabolomics on the two families showing the strongest age-related increase in survival. Our results reveal that ageing in *Magallana gigas* is characterized by coordinated epigenetic, transcriptional, and metabolic reprogramming that reduces host permissiveness to POMS. We show that the epigenetic remodeling of key immune regulators (e.g., Toll-like receptors, MyD88) aligns with transcriptional rewiring of NF-κB and ubiquitin pathways, producing a finely tuned innate immune state marked by enhanced antiviral activity but reduced antibacterial responsiveness. We also identify age-related repression of mTOR signaling, likely promoting autophagy and improving viral control. These regulatory changes are tightly linked to metabolic adjustments, including reduced TCA cycle flux, remodeled nitrogen metabolism, and altered glutathione dynamics, which collectively support a stress-tolerant, energy-conserving phenotype. Together, our findings reveal a fundamental evolutionary trade-off: juveniles prioritize growth at the cost of viral susceptibility, whereas adults invest in cellular maintenance and antiviral preparedness.

## Introduction

Ageing is a progressive and irreversible biological process marked by the gradual deterioration of physiological functions, which over time reduces organismal performance and increases susceptibility to mortality (López-Otín et al., 2013). It is driven by a set of conserved molecular and cellular alterations, including DNA damage, telomere shortening, epigenetic drift, loss of proteostasis, and mitochondrial dysfunction (Mahmoudi & Brunet, 2012). These hallmarks disrupt key processes such as immune regulation, metabolic balance, oxidative stress detoxification, and cell-to-cell communication (López-Otín et al., 2023).

Ageing was mostly studied in vertebrates, but mollusk bivalves are increasingly used as a model given the inter-species variability of maximum life span, ranging from a couple of years for some vagile species (*Argopecten irradians irradians*) to almost 400 years for sessile species (*Artica islandica*) (Abele & Philipp, 2013). In general ageing in bivalves was shown to induce, at the functional level a lowering of the energetic metabolism, a diminishing of the antioxidant defense, an induction of apoptosis, autophagy, and DNA repair mechanisms, and an attenuated stress response, including immune gene expression (Philipp & Abele, 2010; Husmann et al., 2014). The latter predicts that ageing in mollusks should be accompanied by an increasing vulnerability to diseases. For *Magallana gigas*, increased susceptibility with age to the bacterial pathogen *Vibrio aestuarianus* was well documented (Azéma et al., 2017). Surprisingly, this trend is totally reversed in face of the Pacific Oyster Mortality Syndrome (POMS), where individuals show lower susceptibility through time. Indeed, spat and juvenile oysters exhibit remarkably higher mortality rates when compared to older individuals (Petton et al., 2015; Green et al., 2016; Carrasco et al., 2017; Dégremont, 2013).

POMS is a polymicrobial disease initiated by primary infection with the Ostreid herpesvirus type 1 microvariant (OsHV-1 μVar) targeting and disrupting hemocyte physiology, thus impairing host immunity, and facilitating a secondary bacterial infection, including *Vibrio* species (de Lorgeril et al., 2018; Rubio et al., 2019; Delisle et al., 2022; Clerissi et al., 2023; Oyanedel et al., 2023; Kunselman et al., 2024). Given its panzootic status and the huge pressure it exerts on the oyster industry, a better understanding of the factors and associated molecular mechanisms influencing POMS pathogenesis is necessary (Abbadi et al., 2018; Gittenberger et al., 2016; Hwang et al., 2013; Keeling et al., 2014; Martenot et al., 2011; Mortensen et al., 2016; Peeler et al., 2012; Petton et al., 2021; Roque et al., 2012; Segarra et al., 2010). Permissiveness to POMS is defined as the extent to which an oyster allows the infection process initiated by OsHV-1 μVar, including subsequent dysbiosis and opportunistic bacterial proliferation, to progress to mortality. Survival of POMS depends on a complex interplay between host genetic and epigenetic inherited components (Dégremont et al., 2015; Azéma et al., 2017; de Lorgeril et al., 2018; Gutierrez et al., 2020; Divilov et al., 2019, 2021; Gawra et al., 2023; Valdivieso et al., 2025), the influence of temperature (Duperret et al., 2025), and food availability (Pernet et al., 2019). Although age strongly influences permissiveness to POMS in oysters (Dégremont, 2013; Petton et al., 2015; Green et al., 2016; Carrasco et al., 2017), the molecular mechanisms underlying age-dependent reduced susceptibility to POMS remain totally unknown.

Given both the complexity of POMS pathogenesis in oysters and the profound physiological changes occurring during ageing, single-omic approaches would be insufficient to fully capture the regulatory changes underlying the age-dependent shift in disease susceptibility. Multi-omics integration offers an opportunity to simultaneously assess upstream regulatory layers (e.g., epigenetic modifications such as DNA methylation), intermediate molecular responses (e.g., gene expression changes), and downstream functional responses (e.g., metabolic reprogramming). By integrating these complementary biological layers, it becomes possible to identify coordinated molecular signatures and cross-layer interactions that would remain hidden in isolated datasets, providing a more comprehensive and mechanistic understanding of how ageing reshapes the oyster’s susceptibility to POMS. We induced POMS challenge to oysters aged 4, 16, and 28 months, using four biparental families. Two of them displaying the highest survival to POMS gain in function of age were subjected to an integrative multi-omics analysis combining comparative epigenetics, transcriptomics, and metabolomics. Our findings reveal both shared and family-specific molecular trajectories across age, involving key biological processes such as cellular differentiation and antiviral defense. Metabolomic profiling further uncovered an age-related accumulation of tricarboxylic acid (TCA) cycle intermediates, indicative of enhanced energy mobilization in older oysters. Altogether, our multi-layered data highlight molecular signatures underlying the age-driven transition from a disease-permissive to a non-permissive state, offering novel insights into the improved survival of oysters during POMS outbreaks.

## Materials and Methods

### Production of full-sibling families

Four biparental oyster (H2D, F14R, F11N, and F14V) families (*Magallana gigas*) were produced at the Ifremer hatchery in Argenton (France) in 2020. The Spat (F_1_) were transferred and maintained under biosecured conditions at the Ifremer nursery (Bouin, France). Further details about hatchery and nursery steps are described in (de Lorgeril et al., 2018, 2020). At 4-, 16-, and 28-month-old, the offspring were relocated to the Ifremer facilities in Argenton. Oysters were kept at 23°C and fed *ad libitum* (50/50 mixture of *Tisochrysis lutea* and *Skeletonema costatum*, 1,500 µm³/µL) for 15 days for acclimation (Rico-Villa et al., 2006). The pathogen-free status was confirmed by testing for the absence of OsHV-1 µVar DNA and ensuring that *Vibrio sp.* concentrations were below ten colony-forming units per milligram of tissue, as described in (Petton et al., 2015).

### Production of the viral suspension for the experimental infection

A viral suspension of Ostreid herpesvirus 1 µVar (OsHV-1 µVar) was prepared following the protocol from a mix of several variants described by (Schikorski et al., 2011). From moribund oysters, the tissues were dissected, pooled, and homogenized in sterile artificial seawater using a blender under chilled conditions. The homogenate was subjected to sequential centrifugation steps to remove cellular debris (3,000 × g for 10 min followed by 10,000 g for 20 min at 4°C). The resulting supernatant was filtered through 0.45 µm and 0.22 µm pore-size filters to eliminate bacteria and larger particles, yielding a clarified viral suspension. The quantification of genomics units (viral load) of OsHV-1 µVar was performed by real-time quantitative Polymerase Chain Reaction (qPCR) using a Stratagene Mx3005P Real-Time thermocycler. The qPCR reaction volume (20 μL) was: 5 μL DNA (5 ng/µL), 2 µL of each primer at the final concentration of 550 nM (Eurogentec SA), 1 µL of distilled water, and 10 µL of Brilliant III Ultra-Fast SYBR^®^Green PCR Master Mix (Agilent). The virus-specific primer pairs targeted a region of the OsHV-1 µVar genome predicted to a gene that encodes a DNA polymerase catalytic subunit (open reading frame, ORF) 100AY509253: Forward: 5’-ATTGATGATGTGGATAATCTGTG-3’ and Reverse: 5’-GGTAAATACCATTGGTCTTGTTCC-3’ (Webb et al., 2007; Pepin, 2013). The amplification qPCR program consisted of 3 min at 95°C, followed by 40 cycles at 95°C for 5 s and 60°C for 20 s, with a melting temperature curve of the amplicon to verify the specificity of the amplification. Quantification of viral DNA copies was estimated by comparing the observed Cq values to a standard curve of the dimer primer amplification product cloned into the pCR4-TOPO vector. The final concentration was adjusted to 2.2 × 10 genome copies/µL and stored at –80°C.

### Experimental infection

To investigate the effects of ageing on oysters’ permissiveness to POMS, the four families (called recipients) were exposed to the disease through a cohabitation experiment with OsHV-1 µVar-injected donor oysters at ages 4-, 16-, and 28 months (see below), mimicking the natural transmission of POMS (**Figure 1A** and **Table S1**).

**Figure 1.**
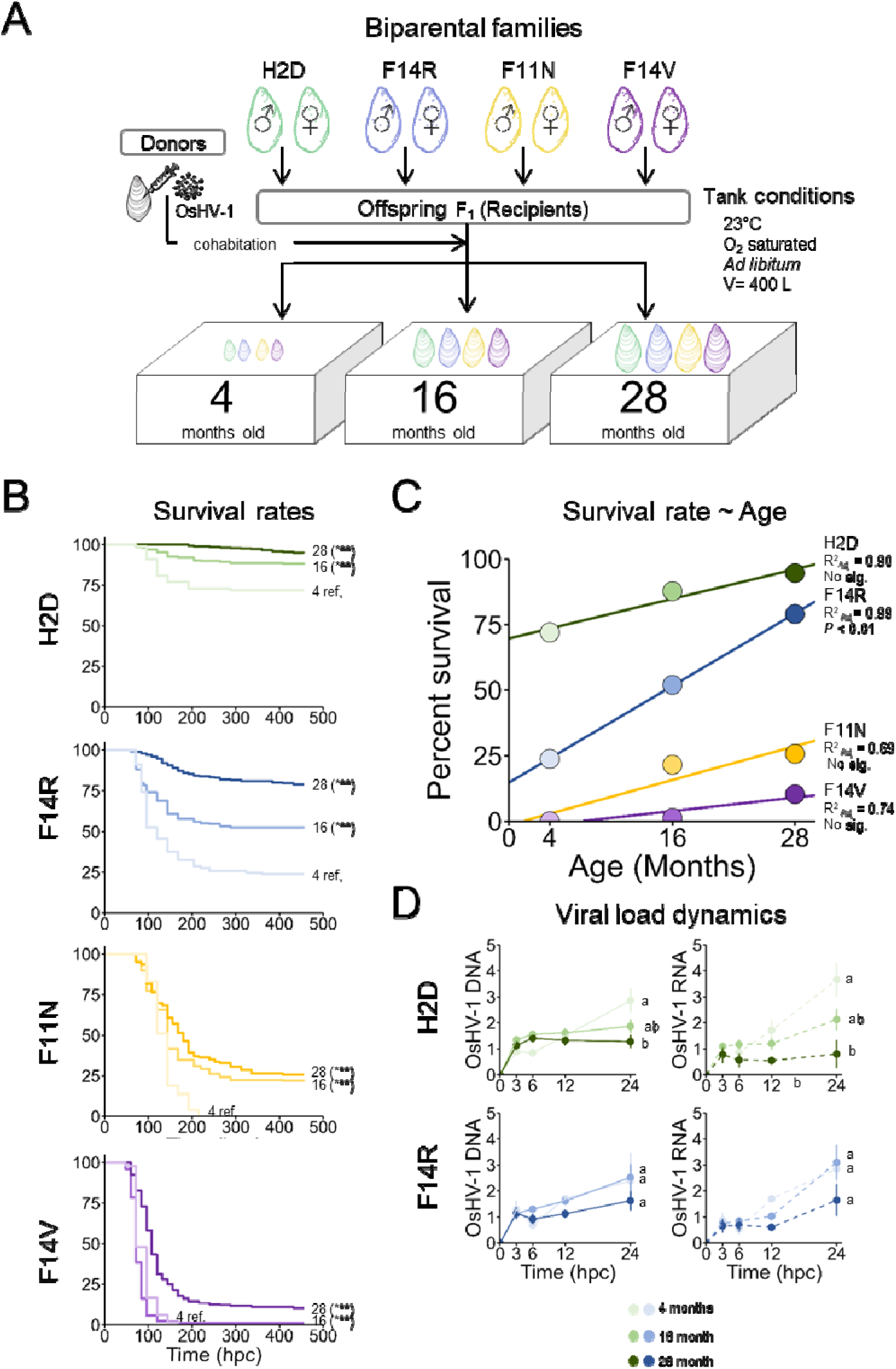
Age-dependent survival and viral load dynamics following OsHV-1 infection for the four oyster biparental families. **A)** Graphical representation of the experimental design of the four biparental Magallana gigas families (H2D, F14R, F11N, F14V) exposed to OsHV-1 μVar through cohabitation with infected donor oysters. The families were challenged at three different ages (4-, 16-, and 28-month-old) under controlled and permissive to the disease tank conditions (23°C, O-saturated, ad libitum feeding, 400 L volume). **B)** Kaplan–Meier survival curves of oyster families H2D, F14R, F11N, and F14V at 4-, 16-, and 28-month-old. Statistical differences were assessed by Mantel-Cox log-rank tests using the 4-month-old group as reference within each family (*** = P < 0.01). **C)** Positive correlations between age and survival rates in all families. **D)** The OsHV-1 μVar DNA load and RNA viral activity kinetics were measured during the first 24 hours post-cohabitation (hpc) in H2D and F14R. Letters denote significant differences among the three age groups at 24 hpc, with the 4-month-old group serving as the reference (One-way ANOVA followed by a Tukey’s post hoc test). Data are presented as the mean ± standard deviation in each group.

Donor oysters came from two stocks of “pathogen-free” oysters to maximize the production of OsHV-1 µVar during cohabitation: one stock, called Dellec, showing a highly susceptible to POMS (> 90%) (de Lorgeril et al., 2018), and one stock, called NSI, with an expected 50–60% mortality (Petton et al., 2013). Twenty hours before the onset of cohabitation, 4-month-old donor oysters were anaesthetized with hexahydrate MgCl_2_ (ACROS, 50 g/L) for 2 hours (Suquet et al., 2009). Then, 100 µL of OsHV-1 µVar suspension was injected into their adductor muscle with a 26-gauge needle using a multi-dispensing pipette (Schikorski et al., 2011). The injected donor oysters were maintained at 23°C in tanks (Volume = 400 L) for 18 hours, an optimal temperature for the disease (Petton et al., 2021). Temperature, pH, salinity, and dissolved oxygen were monitored twice a day with a WTW Multiparameter MultiLine 3630 IDS and a Dotsman P750, equipped with a PT100 sensor and an iButton DS1922L-F5 thermochron.

Cohabitation infection was performed by placing the injected donor and recipient oysters together at a 1:1 biomass ratio. The 4-, 16-, and 28-month-old oysters were acclimated in three, six, and fifteen tanks, respectively, to maintain a constant biomass per liter, as the individual weight increased throughout the age. To increase the transmission of OsHV-1 µVar to the recipient oysters, water renewal was halted for the first 125 hours post-cohabitation (hpc). At the end of this period, all surviving donor oysters were removed from the tanks. Once cohabitation began, six live recipient oysters were collected at 0 (basal conditions), 3, 6, 12, and 24 hours hpc for each family in each experiment. All tissues were removed from the shells, flash-frozen in liquid nitrogen, and stored at –80°C. Mortality was monitored twice daily, and dead oysters were removed as soon as they were observed. The experiment ended when no deaths were recorded in any of the tanks for a continuous 48 hours. The infection experiments were conducted over three consecutive years, each dedicated to a specific age group (4-, 16-, and 28-month-old recipient oysters), under the same conditions and within the same facility with the same donor oysters. To validate the mortality across the three cohabitation experiments, an internal control consisting of 4-month-old NSI recipient oysters was included in each trial.

### DNA and RNA extractions from the same individual oyster

Samples were ground in 50 mL stainless steel bowls with 20 mm grinding balls continuously cooled with liquid nitrogen as described (de Lorgeril et al., 2018) using the Retsch MM400 MILLa device. Grounded samples were stored at –80°C. Following the manufacturer’s protocol, genomic DNA (gDNA) was extracted using the Genomic DNA Tissue Kit (Macherey-Nagel, 740952.250) and stored at −20°C. The gDNA concentration, purity, and quality were assessed using the NanoDrop One spectrophotometer (Thermo Scientific) and 1% agarose gel electrophoresis. Total RNA was extracted using the Direct-zol RNA Miniprep Kit (Zymo Research) according to the manufacturer’s protocol. The RNA concentration was measured using a NanoDrop One spectrophotometer, and integrity was assessed with a BioAnalyzer 2100 (Agilent). All RNA samples were stored at −80°C. To quantify the viral load present in the gDNA samples we followed the same protocol as described above.

### Genome references

For this study, we used the *M. gigas* reference genome (GCA_902806645.1) (Peñaloza et al., 2021; G. Zhang et al., 2012). The *M. gigas* genome comprises a total of 30,418 coding genes, and a description of each gene is provided in **Table S2**. For the virus, we used the OsHV-1 µVar genome (KY242785.1) (Burioli et al., 2017). For this study, we used a previously constructed Gene Ontology (GO-term) gene annotation database (Duperret et al., 2025), needed for the GO_MWU software (see below). Briefly, the coding sequences were extracted with TransDecoder, annotated using ORSON (BLASTX against Uniprot-SwissProt; InterProScan domain prediction), validated with the Omicsbox software (v. 3.4.5) (Gotz et al., 2008), and further enriched with immune-related terms.

### Epigenetic analysis

#### Preparation and library sequencing of samples

DNA methylation libraries were prepared and sequenced by IntegraGen SA (Evry, France) using the NEBNext Enzymatic Methyl-sequencing (EM-seq) protocol. Briefly, 100 ng of extracted genomic DNA, supplemented with unmethylated lambda DNA and methylated puc19 DNA as internal spike-in controls, was used for library preparation. Sequencing was performed on an Illumina NovaSeq platform with 150 bp paired-end reads.

#### Processing and analysis of the samples

The raw paired read quality control for each sample was analysed using FastQC (v.0.53) software (Andrews, 2010), and the adapters were trimmed using TrimGalore! (v.0.6.10) software (Krueger et al., 2023) with: –q 30 –– paired ––clip_R1 5 and ––clip_R2 5 ––Illumina. Any remaining adapters were removed in a second trimming round with default parameters. We used the *’Bismark’* (v0.24.2) software (Krueger & Andrews, 2011), employing the *’bismark_genome_preparation’* function to bisulfite convert the *M. gigas* genome reference. Then, the trimmed reads were aligned to the bisulfite converted *M. gigas* genome using the *’bismark’* function: –q –N 0 –– score_min L,0, −0.4. The duplicated reads were removed using the ′*deduplicate_bismark*′ function, and the methylation extraction calling was accomplished using the ′*bismark_methylation_extractor*′ function: –– no_overlap ––cutoff 8. We applied the same methodology to bacteriophage lambda to assess enzymatic conversion efficiency. We discarded the samples with < 98.0% of conversion efficiency. We used the *’methylKit’* package (v.1.24.0) for data treatment (Akalin et al., 2012), and we retained only the common CpGs present in all samples with a minimum coverage of 8X. For normalization, we used the “median” method. Because the methylated fraction in bivalves primarily occurs within gene body regions (Gavery & Roberts, 2014; Männer et al., 2021; Venkataraman et al., 2020, 2022), we intersect the CpGs located within gene body regions. First, we selected the genic coordinates (chromosome, start, and end positions) of all coding genes in *M. gigas* using the *’biomaRt’* package (v.2.54.1) (Durinck et al., 2009). We then employed the *’foverlaps’* function from the *’data.table’* package (v.1.14.8) (Dowle et al., 2019) to overlap these gene coordinates with the identified CpG coordinates to compile a list of genes containing CpGs within their boundaries. Finally, we calculated the average methylation values for those genes that contained multiple CpG sites within the same gene body.

### Transcriptomic analysis

#### Preparation and library sequencing

RNA-seq libraries were prepared using the NEBNext Ultra II Directional RNA Library Prep Kit. Paired-End (PE) reads sequencing was performed with 100 base pair reads on the Illumina NovaSeq sequencer. Library construction and sequencing were carried out by IntegraGen SA (Evry, France)

#### Processing and analysis of samples

The raw paired reads were trimmed using Trim Galore! software (v.0.6.7) (Martin, 2011; Krueger et al., 2023) with parameters: –q 30 ––illumina ––stringency 1 –e 0.1 ––length 35. The quality of the sequences was assessed before and after trimming using FastQC software (v.0.11.9) (Andrews, 2010). We mapped the trimmed PE reads to the *M. gigas* genome using the following parameters: ––runMode alignReads, ––outSAMmapqUnique 60, and ––outSAMattributes All with the STAR software (v.2.7.10b) (Dobin et al., 2013). Finally, count reads were obtained using the HTSeq software (v.0.9.1) (Putri et al., 2022) with: ––stranded reverse –t gene –i ID –m union –q ––minequal 10. The trimmed reads were also mapped to the OsHV-1 µVar with Bowtie2 software (v2.3.4.3) (Langmead et al., 2009; Langmead & Salzberg, 2012) to analyze the viral transcriptional activity. The HTSeq-count software was used to obtain the number of counts per ORF with the same parameters as above, except for the feature type as CDS and the ID attribute as product. The viral raw counts were normalized based on the formula: 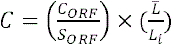. While the first ratio corrects for the size of the ORF by dividing the raw counts (*C_ORF_*) with the size of it (*S_ORF_*), the second one corrects the sequencing depth with a ratio of the average reads for all the time points 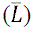 and the number of total reads for the specific time points (*L_i_*), as described (de Lorgeril et al., 2018). We used the *’DESeq2’* package (v.1.42.1) for data treatment (Love et al., 2014), and raw count gene matrix was filtered (counts ≤ 15 across all the samples), and library normalization was applied, followed by a variance-stabilizing transformation with the *’vst’* function.

#### Metabolomic analysis

The Montpellier Alliance for Metabolomics and Metabolism Analysis (MAMMA) platform (BioCampus, Montpellier, France) carried out targeted metabolites on global metabolites (GM), nucleotide metabolites, and metabolites involved in the tricarboxylic acid (TCA) cycle. Briefly, 20 mg of powder samples were supplemented with 980 µL of ice-cold methanol (0013684102BS, Biosolve), 5 µL of 2-(N-Morpholino)-ethanesulfonic acid (M8250, Sigma-Aldrich) and 20 µL of mixed internal standards along with 0.5 g of glass beads. Samples were purified via SPE column and dried in a Speedvac (Turtoi, Jeudy, Henry, et al., 2023).

#### Processing and analysis of the samples

For the GM analysis (Turtoi, Jeudy, Valette, et al., 2023), samples were reconstituted in 200 µL of UPLC water and transferred to LC-MS polypropylene vials. UHPLC 1290 Infinity II with an HPLC Discovery^®^ HS F5, 3 μm particle size, L × I.D. 15 cm × 2.1 mm (567503-U, Sigma-Aldrich) was used for chromatographic analysis. GM profiling analysis was performed on a 6495 LC/TQ mass system and the Agilent MassHunter workstation (Agilent, Santa Clara, USA) with the dynamic MRM scan mode. The nucleotide metabolites analysis was conducted using chromatography on a UHPLC 1290 Infinity II system equipped with an InfinityLab Poroshell HPH-C18 column (2.7 μm particle size; L × I.D. 100 cm × 3.1 mm; 695975-502, Agilent). The mobile phase A consisted of 10 mM ammonium acetate (5.33004; Sigma-Aldrich), 2.5 μM medronic acid (5191-3940; Agilent), and 0.1% formic acid in UPLC-grade water (0023214102BS; Biosolve). The mobile phase B consisted of 90% UPLC-grade methanol (0013684102BS; Biosolve), 10% UPLC-grade water, 2.5 μM medronic acid, and 0.1% formic acid. The gradient for the analysis was as follows: 2% B at 0 min; 2% B at 1 min; 42% B at 4 min; 100% B at 5 min; 100% B at 6 min; 2% B at 6.5 min; and 0% B at 8 min. The column temperature was maintained at 30°C, with a flow rate of 0.4 mL/min and an injection volume of 3 μL per run. Nucleotide metabolite profiling was performed using a 6495 LC/TQ mass spectrometer utilizing the dynamic MRM scan mode. Source parameter settings for both positive and negative ion modes were as follows: capillary voltage at 3.5 kV, gas temperature at 120°C, gas flow at 11 L/min, nebulizer pressure at 40 psi, sheath gas temperature at 400°C, and sheath gas flow at 12 L/min. The Krebs cycle metabolites analysis was performed using gas chromatography coupled with mass spectrometry (GC-MS). Purified and dried samples were reconstituted in 200 µL of an 80:20 UPLC methanol/UPLC water solution, and 10 µL of d27-Myristic acid was added as an internal standard (75 µg/mL; 400505, Agilent). The samples were then transferred into brown crimp vials with inserts and dried using a SpeedVac. Derivatization was performed by a PAL autosampler as follows: 90 µL of methoxyamine hydrochloride was added to the samples (20 g/L; 89803, Sigma-Aldrich), which were incubated at 30°C for 90 min. Subsequently, 10 µL of MSTFA (TS-48915; Thermo Scientific) was added, followed by incubation at 37°C for 30 min. Chromatographic analysis was performed on an Agilent 8890 GC system equipped with a DB-MS + DG column (0.25 μm film thickness, L × I.D. 30 m × 0.250 mm; 122-5532G, Agilent). One microliter of the prepared sample was injected in split mode (10:1), with the inlet temperature maintained at 250°C. Helium was used as the carrier gas at a flow rate of 1.1 mL/min. The GC temperature program started at 60°C for 1 min, followed by a ramp of 60°C/min to 120°C, then 30°C/min to 250°C, and finally 60°C/min to 325°C, with a 5-min hold at the final temperature. Krebs cycle metabolite profiling was conducted using a 5977B MSD mass spectrometer with dynamic MRM scan mode. The temperature was set to 250°C, and the electron energy was maintained at 70 eV. The peak area counts were normalized based on their respective internal standard control compounds for each detected metabolite from the GM, Nucleotides, and Krebs cycle. The *’MetaboAnalystR’* package (v.4.0.0) (Chong & Xia, 2018; Chong et al., 2019; Pang et al., 2020) was used to filter compounds based on an interquartile range (IQR) cutoff of < 25, removing 10% of the data. Subsequently, quantile normalization was applied, followed by a logarithmic transformation.

#### Statistical analyses

Statistical analyses were performed using R software (v.4.4.1). Plots were primarily generated using the *’ggplot2’* package (v.3.5.1) (Wickham, 2009) unless otherwise. To ensure data quality, samples falling outside the 95% confidence ellipse in PCA were considered outliers and subsequently removed.

#### Survival curves

Survival curves (Kaplan-Meier model) were generated using the *’survfit’* and plotted using the *’ggsurvplot’* functions from the *’survival’* (v.3.2-11) (Therneau & Lumley, 2015) and the *’survminer’* (v.0.4.9) packages (Kassambara et al., 2017), respectively. The Cox proportional hazards model was performed using the *’coxph’* function from the *’survival’* package.

#### Viral DNA and RNA load in oyster

Normality and homogeneity of variance for viral DNA loads and RNA transcription were evaluated using the functions of *’shapiro*.*test’* from the *’stats’* (v.4.4.1) and *’levenetest’* from the *’car’* packages (Fox & Weisberg, 2019), respectively. A one-way ANOVA was conducted with the *’aov’* function to compare viral DNA and RNA loads across age groups, followed by Tukey’s Honest Significant Difference (HSD) test using the *’Tukeyhsd’* function from *Stats’* package.

#### Weighted Correlation Network Analysis

To identify biologically relevant co-expression modules associated with age, we applied a Weighted Correlation Network Analysis (WGCNA) using the *’WGCNA’* package (v.1.73) (Langfelder & Horvath, 2008). This analysis was performed across the three omics layers per family: epigenetics (DNA methylation), transcriptomics (gene expression), and metabolomics (metabolites). For all the omics, the trait of interest was the age, and samples were re-coded as follows: 4 months =1, 16 months = 2, and 28 months = 3. The associations between the network modules and age were assessed using the Pearson (r) correlation (*P* < 0.05). For each dataset, an unsupervised co-expression network was constructed using the following parameters: maxBlockSize = 30,000, networkType = signed, TOMtype = signed, mergeCutHeight = 0.25, and minModuleSize = 100. A soft power threshold was selected to achieve a scale-free topology, optimizing the identification of biologically meaningful network structures. Modules showing either significant positive or negative correlations with age were retained, and when multiple modules shared the same direction, they were merged to generate a unified list of age-associated positive and negative features (i.e., gene methylation, gene expression and metabolites). To evaluate the biological significance of age-associated modules in epigenetic and transcriptomic analyses, we retrieved the kME values (eigengene connectivity) of all genes present in the significant modules, considering both positive and negative associations. The analysis included all 30,418 coding genes, assigning kME values to the genes retrieved from the significant modules, while giving a value of 0 for the genes outside the modules. We use the GO_MWU package (github.com/z0on/GO_MWU) (Wright et al., 2015), focusing on the enrichment analysis of Biological Process (BP) GO-terms with the following parameters: largest = 0.5, smallest = 10, clusterCutHeight = 0.25, Module = TRUE, and Alternative = g. The BP GO-terms were significant at *P.adj* < 0.05. To assess the impact of age on metabolomics, we used the normalized metabolite data from the *’MetaboAnalystR’* package as input for WGCNA analysis, identifying significant modules associated with age. To simplify and summarize the list of significant BP GO-terms obtained in epigenetics and transcriptomics from GO_MWU, we categorized the BP GO-terms into broader groups based on the Gene Ontology database (**Table S3**). This grouping, based on the current understanding of oyster–POMS interactions, further supports the rationale for organizing these categories according to established knowledge of POMS pathogenesis (de Lorgeril et al., 2018, 2020; Duperret et al., 2025).

The metabolites within the significant modules associated with age were further used as an input list for pathway reconstruction and enrichment analysis using the *’FELLA’* package (v.1.22.0) (Picart-Armada et al., 2018), based on a diffusion algorithm. Specifically, each input metabolite introduced a unitary flow model that propagated through intermediate KEGG entities (including reactions, enzymes, and modules) to identify biological pathways in which the metabolite is involved. Once identified, we extracted the genes annotated to participate in the significantly enriched KEGG pathways by querying KEGG through the ‘KEGGREST’ package (v.1.46.0) (Tenenbaum, 2017) with database *M. gigas* (T03920), thereby enabling the integration of metabolomic findings with the corresponding genes.

## Results

### Ageing reduces viral replication in POMS-susceptible oyster families

Following the three infection challenges (one for each age: 4-, 16-, and 28-month-old), mortality decreased with ageing for the four families (**Figures 1B**–**C**). For each infection challenge, both the 4-month-old NSI donors and the 4-month-old NSI recipient oysters (positive control) deployed in the experimental tanks showed similar survival curves across the three experiments (**Figure S1A**–**B**), confirming that the mortality patterns observed in the four families tested were not biased by differences in experimental infections (virus load or infectious efficiency). Only the F14R and H2D were selected for further analysis due to their contrasting age related gains of survival (**Figure 1B**). Key infection indicators revealed that 28-month-old oysters in H2D showed significantly lower viral DNA loads (One-way ANOVA: F□,□□= 5.9, *P* < 0.05) and reduced viral transcriptional activity (F□,□□= 8.2, *P* < 0.01) compared with their 4□month old siblings (**Figure 1D**). Whereas age-related differences are not significant in F14R (viral DNA: F□,□□= 0.74, *P* = 0.50; and viral transcription: F□,□□= 1.4, *P* = 0.28), although a trend toward a decrease in viral load and transcriptional activity was apparent in the oldest oysters (**Figure 1D**). Overall, our findings demonstrated that ageing limits OsHV-1 µVar replication associated with increased survival. To investigate the molecular mechanisms underlying age-related changes influencing permissiveness to POMS, we analyzed the methylome, transcriptome, and metabolome of 4-, 16-, and 28-month-old oysters just before the infection challenge (0 hpc)

In the epigenetic dataset, each sample yielded an average of 84,991,198□±□8,765,131 trimmed reads, with mapping of 42.87□±□2.52% and a duplicate read rate of 15.20 ± 3.35%. The enzymatic conversion efficiency estimated by the lambda genome spike-in positive control was 99.18% ± 3.62% (**Table S4**). We identified 2,164,482 CpG sites in F14R and 2,300,773 CpG sites in H2D that were shared across the three age groups in each family. Both families exhibited age-related methylation patterns (**Figures S2A** and **S2B**). For further analysis, we averaged the methylation values at the gene level (within each gene’s body region) for each sample and age group, resulting in two gene-level methylation lists (26,104 genes for F14R and 26,534 genes for H2D, **Table S5** and **Table S6**, respectively). For the transcriptomic data, each sample yielded an average of 86.65% ± 1.32% alignment rate (**Table S4**) and showed a clear age-dependent pattern of gene expression in both families (**Figures S2C** and **S2D**). For the metabolomic data, based on 76 filtered out of the initial 114 metabolites, a strong separation between 4-month-old and older oysters (16– and 28-month-old) was observed, highlighting a remarkable age-related metabolic shift (**Figures S2E** and **S2F**). Our results revealed that ageing induced profound changes in methylome, gene expression, and metabolism, which could govern physiological changes influencing permissiveness to POMS. Outlier removal resulted in the exclusion of two epigenetic, one transcriptomic, and four metabolomic samples (**Table S4**).

### Ageing drives a shift from a permissive to a non-permissive state to POMS, reshaping critical pathways that enhance the host’s ability to cope with POMS

Ageing drives a shift from a permissive to a non-permissive state to POMS, reshaping critical pathways that enhance the host’s ability to cope with infection. To investigate the molecular basis of this age-dependent shift, we sought to integrate epigenetic, transcriptomic, and metabolomic layers. Since changes in DNA methylation, gene expression, and metabolite profiles are expected to be coordinated and functionally linked, we applied a Weighted Gene Co-methylation/Co-expression/Co-metabolite Network Analysis. This systems-level approach allowed us to identify age-associated molecular modules significantly correlated (positively or negatively) with age. To further characterize the biological meaning of these modules, we performed enrichment analyses of BP GO-terms, revealing functional pathways that either converge or diverge between families.

### Epigenetics reflects age-dependent changes in immunity

At the epigenetic level for F14R, we identified one positively correlated module (693 methylation genes; r = 0.73; *P* < 0.001) and two negatively correlated modules (574 and 425 methylation genes; r = −0.78 and −0.50; *P* < 0.001 and 0.05, respectively) with age (**Figure S3A** and **Table S7**). For H2D, we identified two positively correlated modules (660 and 163 methylation genes; r = 0.55 and 0.63; both *P* < 0.05) and one negatively correlated module (2,085 methylation genes; r = −0.86; *P* < 0.001) with age (**Figure S3B** and **Table S7**). From these gene methylation sets, 10 were overrepresented and 9 underrepresented GO-terms for the F14R, while it was 10 and 23 GO terms for the H2D, respectively (**Table S8**). The BP GO-term analysis revealed biological processes that are either common or specific to each family. For both families, genes with increased DNA methylation levels with age were enriched in biological regulation, developmental processes, responses to stimuli, and immune system processes. Among the genes showing decreased DNA methylation levels with age, the BP GO-terms were mostly related to cellular processes and localization. Divergent epigenetic signatures mainly concerned the H2D, where several biological processes linked to biological regulation, developmental processes, and reproduction were enriched among genes with decreased DNA methylation levels, while processes related to localization were enriched among genes where DNA methylation levels increased with age (**Figure 2A**−**2B**). Overall, these results aligned with biological expectations of ageing in bivalves, showing a shift from cellular proliferation and structural modification in younger oysters to a remodeling of metabolism, stress responses, and immune functions in the older ones.

**Figure 2.**
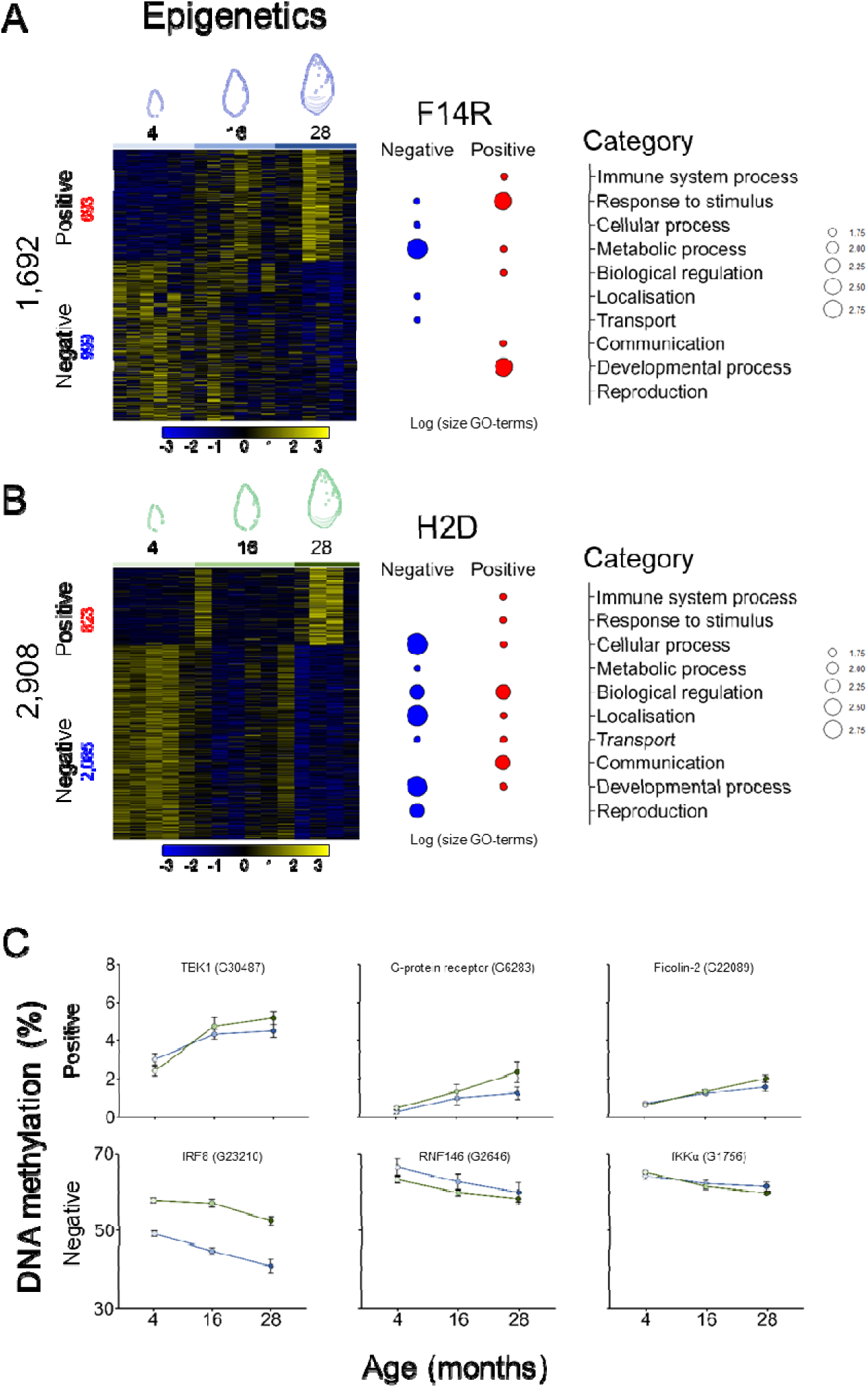
Age-associated DNA methylation modules and Gene Ontology enrichment (Epigenetics) Heatmaps of age-associated DNA methylation (Epigenetics) Heatmaps displaying genes with positive (red) or negative (blue) associations between DNA methylation and age, alongside a summary of enriched Gene Ontology (GO) terms of the Biological Process category for **A)** F14R and **B)** H2D. Complete GO-term lists are available in Table S6. **C)** Examples of immune/stress genes showing concordant age-associated gene body methylation rate trends in both families (F14R, blue; H2D, green).

Given the pivotal role of immune and stress responses in host-pathogen interactions in oysters, we focused on the genes belonging to the Response to stimulus and Immune system process GO-terms. Between the 693 and 823 positively, and the 999 and 2,085 negatively age-associated methylation genes in F14R and H2D, respectively (**Figures S3A**–**3B**), a total of 19 (13 + 6) immune– and stress-related genes were shared (**Figure S4A**). These shared genes were involved in pathogen recognitions, such as the C1q domain-containing protein (G18427), complement C1q tumor necrosis factor-related protein 2 (G3352), and ficolin-2 (G22089); in immune signaling as the inhibitor of nuclear factor kappa-B kinase subunit α (IKKα, G1756), serine/threonine-protein kinase H1 homolog (G20959), interferon regulatory factor 8 (IRF8, G23210), TNF receptor-associated factor 2 (TRAF2, G25366), serine/threonine-protein kinase TBK1 (G30487); and in immune regulation as the E3 ubiquitin ligases TRIM63 (G10197), TRIM45 (G22126), TRIM56 (G25021), tripartite motif-containing protein 2 (G15281), RNF146 (G2646), FMRFamide receptor (G3282), interleukin-6 receptor subunit β (G628), G-protein coupled receptor family 1 domain-containing protein (G6283) B-cell lymphoma/leukemia 11A BCL11A (**Figure S4A**). Among the shared genes, we examined the IRF8 (G23210), TBK1 (G30487), RNF146 (G2646), G-protein coupled receptor (G6283), IKKα (G1756), and ficolin-2 (G22089), and all genes displayed aligned age-associated DNA methylation trends (increases or decreases) in both families (**Figure 2C**). Additionally, we detected Toll-like receptors (TLRs), critical components of innate immune recognition, which also exhibited notable age-associated methylation patterns, such as the toll-like receptor 4 (G26336) in F14R and the toll-like receptor 4 (G9660 and G9701) and toll-like receptor 13 (G21066) in H2D. This family-specific modulation of TLR at the methylation level may reflect nuanced changes in pathogen sensing and immune responsiveness across the lifespan, underscoring the complexity of immune epigenetic regulation in oyster host-pathogen interactions. Overall, these results indicated that age-associated DNA methylation changes also occur in immune– and stress-related genes, suggesting that epigenetic regulation at the methylome level during ageing contributes to the modulation of the immune landscape in oysters. Such epigenetic regulation may also influence the capacity of oysters to respond to OsHV-1 µVar infection, potentially explaining why the older oysters displayed reduced permissiveness to POMS compared to the younger ones. To investigate whether these age-related epigenetic modifications are also reflected at the transcriptional level, either gene-to-gene or biological functions, we analyzed the gene expression patterns across the same age groups.

### Ageing-associated transcriptomic shifts reflect metabolic and immunity remodeling

At the transcriptomic level in F14R, we identified 14,546 genes grouped into six modules correlated with age: two positively (6,468 and 1,420 genes; r = 0.96 and 0.50; *P* < 0.001 and 0.05, respectively) and four negatively (4,517; 1,380; 463; and 298 genes; r = −0.96, −0.70, −0.80, and −0.55; *P* < 0.001, 0.01, 0.001, and 0.05, respectively) modules (**Figure S3C** and **Table S9**). In the H2D, we identified 11,533 genes grouped into nine modules correlated with age: five positive (2,253; 1,975; 1,558; 313; and 214 genes; r = 0.93, 0.75, 0.76, 0.63, and 0.51; *P* < 0.001, 0.001, 0.001, 0.01, and 0.05, respectively) and four negative modules (3,184; 1,771; 133; and 105 genes; r = −0.91, −0.91, −0.49, and −0.53; *P* < 0.001, 0.001, 0.05, and 0.05, respectively) (**Figure S3D** and **Table S9**). The enrichment analysis revealed 167 positively and 175 negatively BP GO-terms in F14R, and 95 positively and 195 negatively BP GO-terms in H2D correlated with age (**Table S10**). Among these, 181 BP GO-terms (77 positive and 104 negative) were in common (**Table S11**). To synthesize the important number of BP GO-terms, as we did in epigenetics, we classified the BP GO-terms from transcriptomic analysis into broader functional categories. Among the significant BP GO-terms, those most directly involved between ageing and susceptibility to POMS included over-representation pathways in the Immune system process (e.g., Cell surface receptor signaling pathway and Defense response to symbiont) and antiviral immunity (e.g., Defense response to virus), and to the under-representation of Toll-like receptors and Myd88 (e.g., Toll signaling, and MyD88-dependent toll-like receptor signaling pathways), along with regulation of canonical NF-kappaB signal transduction. DNA repair mechanisms (e.g., DNA damage response and Regulation of DNA repair) were exclusively positively associated with age, suggesting a conserved activation of these protective pathways across both families. The response to stimulus and homeostatic processes was in both positive and negative sets of genes; however, the results were quite nonspecific among the genes positively associated with age (e.g., Cellular response to stimulus). A strong and specific under-expression of genes in the response to oxidative stress and the maintenance of homeostasis processes was also evidenced (e.g., Response to oxidative stress, the Wnt signaling, and Cellular homeostasis). Additionally, the enrichment analysis underscored the regulation of a broad spectrum of metabolic processes. Among the genes positively associated with ageing, those involved in protein catabolism (e.g., Proteolysis involved in protein catabolic process) and nucleotide metabolism (e.g., Nucleobase-containing compound metabolic process) were evident. For the genes negatively associated with age, the metabolism of protein processes was shown to be under-represented (e.g., Proteolysis) as well as the metabolism of lipids (e.g., Lipid metabolic process). Between families, differences in metabolic processes were more pronounced in H2D, which showed a marked under-expression of genes involved in energy production (e.g., ATP biosynthetic process, Generation of precursor metabolites and energy, and Energy derivation by oxidation of organic compounds) (**Figure 3A**−**3B**).

**Figure 3.**
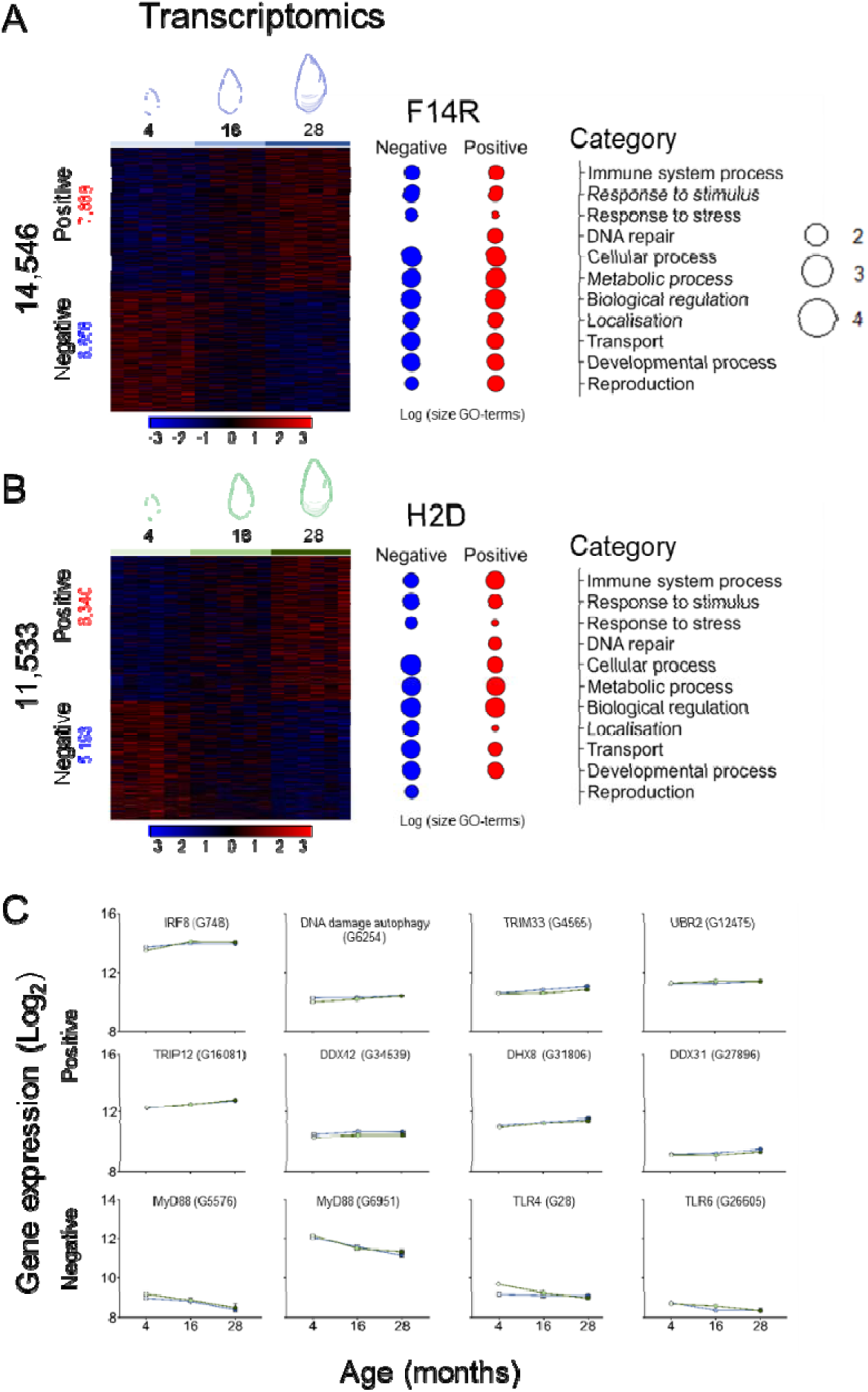
Age-associated gene-expression modules and GO enrichment (Transcriptomics) Heatmaps of age-associated gene expression (Transcriptomics). Heatmaps displaying genes with positive (red) or negative (blue) associations between gene expression and age, alongside a summary of enriched Gene Ontology (GO) terms of the Biological Process category for **A)** F14R and **B)** H2D. Complete GO-term lists are available in Table S7. **C)** Examples of immune/stress genes showing concordant age-associated gene expression (Log2 of normalized counts (VST)) trends in both families (F14R, blue; H2D, green).

We also extended our analysis to the genes exhibiting transcriptional regulation involved in Response to stimulus and Immune system process GO-terms in the context of POMS. Between the 7,888 and 6,340 positively, and the 6,658 and 5,193 genes negatively age-associated in F14R and H2D (**Figures S3C**−**D**), 174 (150 + 24) immune– and stress-related genes were shared (**Figure S4B** and **Table S12**). Specifically, we found age-associated increases in the expression of 150 genes, including the interferon regulatory factor 1 (IRF8, G748), DNA damage-regulated autophagy modulator protein 2 (G6254), a set involving E3 ubiquitin-protein ligases genes such as (TRIM33, G4565), (UBR2, G12475), and (TRIP12, G16081) which are key regulators of protein turnover and immune signaling, and ATP-dependent RNA and DNA helicases genes including (DDX42, G34539), (DHX8, G31806) and (DDX31, G27896) genes, which are essential for RNA metabolism, DNA repair, and antiviral defense (**Figure 3C**). On the other hand, we observed an age-associated decrease in the expression of 24 genes in innate immune signaling, including multiple genes of myeloid differentiation primary response protein MyD88 (e.g., G6951 and G5576) and related to Toll-like receptor (TLR) pathways such as TLR 4 (G28) and TLR 6 (G26605) (**Figure 3C**). Overall, these results indicated that ageing in oysters orchestrates a complex and coordinated biological program, marked by enhanced DNA repair capacity, a reshaped immune landscape with heightened antiviral response genes, and reduced NF-κB signaling. In the context of POMS, such gene expression reprogramming may promote the improved survival of older oysters by strengthening intracellular antiviral defenses and repair mechanisms, while tempering the excessive inflammatory stress responses. Consistent with previous observations in ageing in bivalves (Abele et al., 2009), we also detected a decline in energetic metabolism and antioxidant defenses. Given the critical role of energy metabolism in determining susceptibility to POMS in oysters, we next performed a metabolomic analysis across the same age groups.

### Global metabolomic analysis reveals age-dependent decline in mitochondrial function and central metabolism

At the metabolomic level in F14R, 21 metabolites were clustered into a module positively correlated (r = 0.94, *P* < 0.001), while 35 metabolites were in two modules negatively correlated with age (r = −0.90 and −0.76, *P* < 0.001) (**Figure S3E**). In H2D, 23 and 26 metabolites were grouped into one positive and one negative module (r = 0.85 and −0.92, *P* < 0.001, respectively) correlated with age (**Figure S3F**). The most pronounced differences were between the 4-month-old oysters and the older age groups (16– and 28-month-old) (**Figure 4**), where a metabolic shift was evident in key amino acids (e.g., alanine, creatine, cysteine, argininosuccinic acid) and nucleotides (e.g., AMP, GMP, UMP), as well as in energy-related metabolites (e.g., NAD). Notably, a progressive decline was observed in several TCA cycle intermediates, such as fumarate, malate, and citrate, indicating a reduction in mitochondrial oxidative metabolism with age. Consistent with transcriptomic data, this metabolic shift was more pronounced in H2D. A set of 37 (15 + 22) metabolites was shared between the two families, including central metabolic intermediates involved in ATP generation, redox homeostasis (e.g., glutathione derivatives), nucleotide turnover, and amino acid metabolism (**Figure S5**). These patterns revealed a conserved metabolic aging signature characterized by mitochondrial downregulation, altered redox balance, and remodeling of amino acid and nucleotide metabolism. To gain deeper functional insights into the metabolic reprogramming associated with ageing in oysters, we performed enrichment analysis of the positively and negatively age-associated metabolites using the *’FELLA’* package. This approach integrates metabolomic profiles with the *M*. *gigas* KEGG database, enabling the identification of enriched KEGG pathways. Results revealed family-specific KEGG pathways, such as the Toll-like receptor signaling pathway in F14R (**Figure S6A**) and Pyruvate metabolism in H2D (**Figure S6B**). To obtain a broader view of metabolism and ageing in oysters, we analyzed the 37 common age-associated metabolites shared by both families. This analysis identified KEGG pathways, including those involved in amino acid metabolism, carbohydrate metabolism (e.g., the TCA cycle), and nucleotide metabolism. Pathways related to genetic information processing, such as RNA degradation, protein export, and mRNA surveillance, were also enriched. Notably, we observed enrichment of cellular processes previously linked to POMS outcomes, including autophagy, mitophagy, peroxisome activity, and the mTOR signaling pathway, all of which are central to antiviral defense, oxidative stress control, and energy homeostasis in oysters. Regulation of neuroactive ligand–receptor interaction and related signaling pathways further suggests an interplay between metabolic processes and neuroendocrine regulation, which may modulate immune and stress response (**Figure S6C**). Together, these findings highlighted a core set of metabolic and regulatory networks potentially underpinning age-dependent permissiveness to POMS.

**Figure 4.**
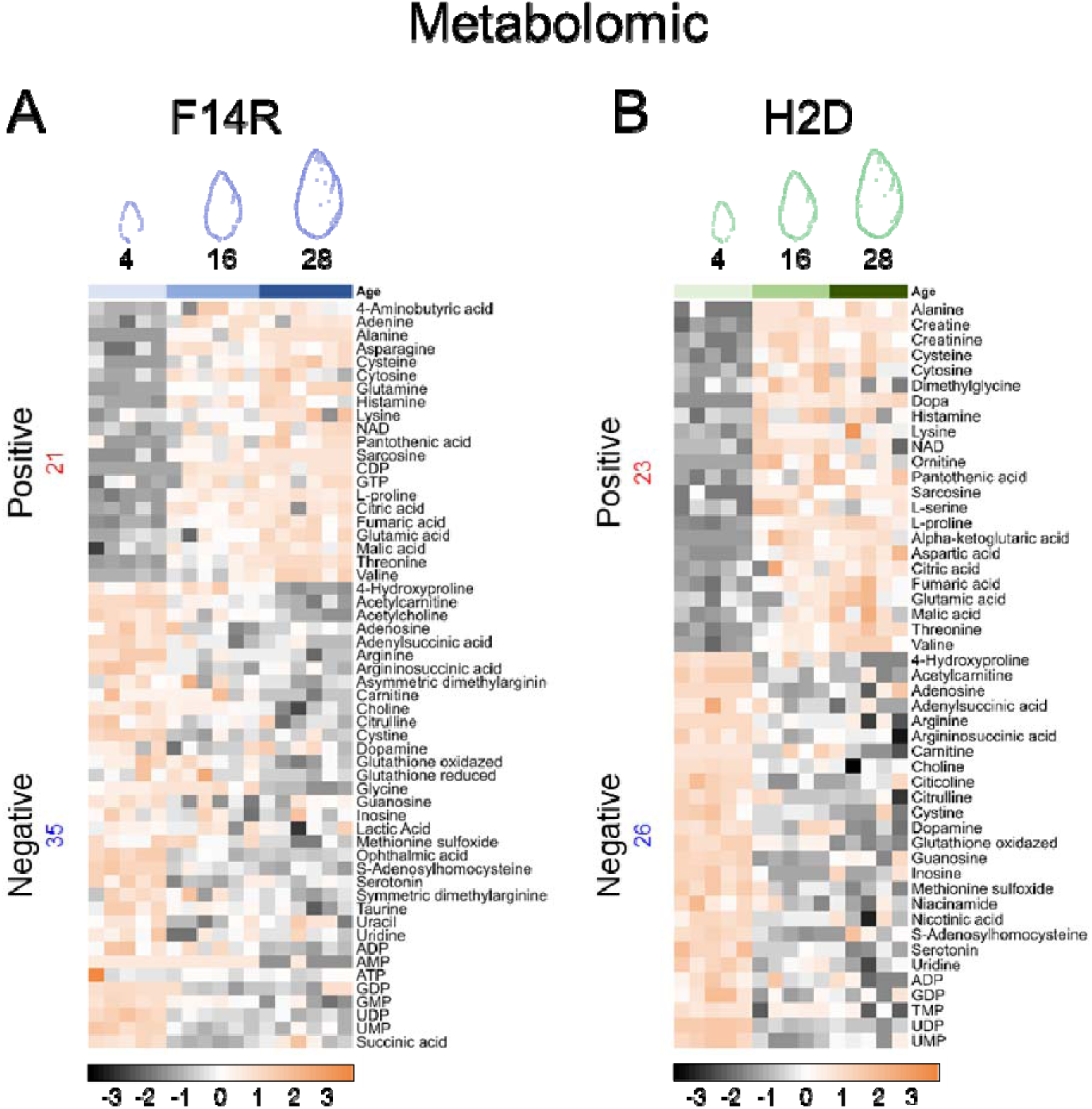
Age-associated metabolites (Metabolomics). Heatmaps displaying metabolites with positive (red) or negative (blue) associations with age for **A)** F14R and **B)** H2D.

### Integration of multi-omics: linking epigenomics, transcriptomics, and metabolomics with signals of ageing influencing permissiveness to POMS

#### Epigenetic and transcriptomic relationship in age-dependent modulation of immune and stress responses

To explore the relationship between epigenomic and transcriptomic layers, we overlapped the list of genes showing age-related methylation changes with those exhibiting age-associated expression patterns for both positive and negative associations. This gene-to-gene overlap between epigenomic and transcriptomic lists revealed 855 genes in F14R and 1,282 genes in H2D, but only 74 were in common between the two families. Although we identified the same set of genes associated with age at both the epigenetic and transcriptomic levels, it has been consistently shown in invertebrates that DNA methylation levels do not directly correlate with gene expression levels (Dixon & Matz, 2022). It seems more plausible that DNA methylation interacts with other components of the epigenetic machinery, such as epigenetic writers (Histone modifying enzymes) or readers (Methyl binding proteins) that influence the chromatin accessibility, and there exists a more complex interrelationship with epigenetics and transcription regulation in invertebrates (Bogan et al., 2023; G. Wang & Mai, 2025). Moreover, genetic backgrounds may differ between families, and regulatory mechanisms can vary accordingly; nevertheless, these differences often converge on similar biological functions by mobilizing different members of multigenic families (de Lorgeril et al., 2020). To characterize the relationship between age-associated changes in DNA methylation and the gene expression at the biological function and the context of POMS, we focused on the significant BP GO-terms belonging to Response to stimulus and Immune system process in both omics. This targeted approach provided an integrated view of immune– and stress-related processes potentially modulating the permissiveness to POMS in oysters in ageing. Notably, several biological processes, including Defense response to virus, Response to symbiont, Response to oxidative stress, and Chemokine-mediated signaling, displayed consistent directionality for both omics layers in both families, indicating the existence of a core of age-modulated pathways (**Figure 5**). The recurrence of the same biological functions in two independent genetic backgrounds reinforces their potential roles as fundamental components of the immune landscape in oysters. Together, these observations supported the hypothesis that age-related epigenetic modifications may act as upstream regulatory layers, either promoting or repressing transcriptional programs that shape the organism’s permissiveness to POMS.

**Figure 5.**
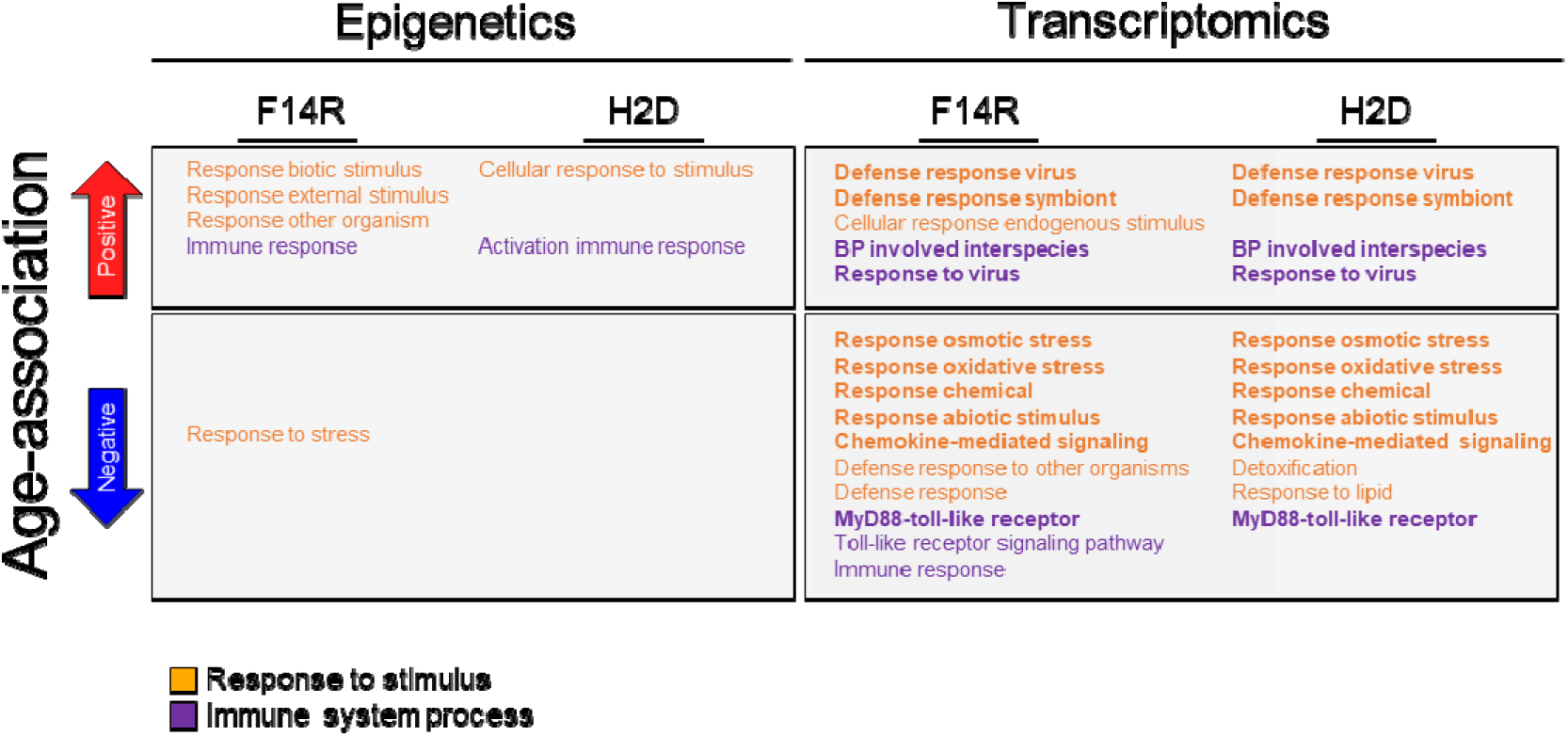
Gene Ontology (GO-terms) enrichment of age-associated immune and stimulus-response genes. Biological Process GO terms related to Response to stimulus and Immune system process, derived from genes with positive (red) or negative (blue) associations with age. Results are presented separately for DNA methylation (epigenomic) and gene expression (transcriptomic) datasets in each family (F14R and H2D).

#### Coordinated age-related metabolic shifts highlight the regulation of oxidative stress and redox imbalance

To validate the functional relevance of the enriched KEGG pathways identified through metabolomics, we compiled all genes annotated to these pathways and cross-referenced them with our transcriptomic datasets to determine age-dependent expression dynamics. Given the large number of KEGG pathways, we focused our validation on seven key pathways: Carbohydrate metabolism (citrate cycle, TCA cycle; crg00020), Nucleotide metabolism (purine metabolism, crg00230; and pyrimidine metabolism, crg00240), Amino acid metabolism (alanine, aspartate, and glutamate metabolism, crg00250; and arginine and proline metabolism, crg00330), Metabolism of other amino acids (taurine and hypotaurine metabolism, crg00430), and Glutathione metabolism (crg00480). These pathways have been linked to POMS permissiveness in relation to temperature, the major driver of the disease (Duperret et al., 2025) (**Figures 6**, **S7,** and **S8**).

**Figure 6.**
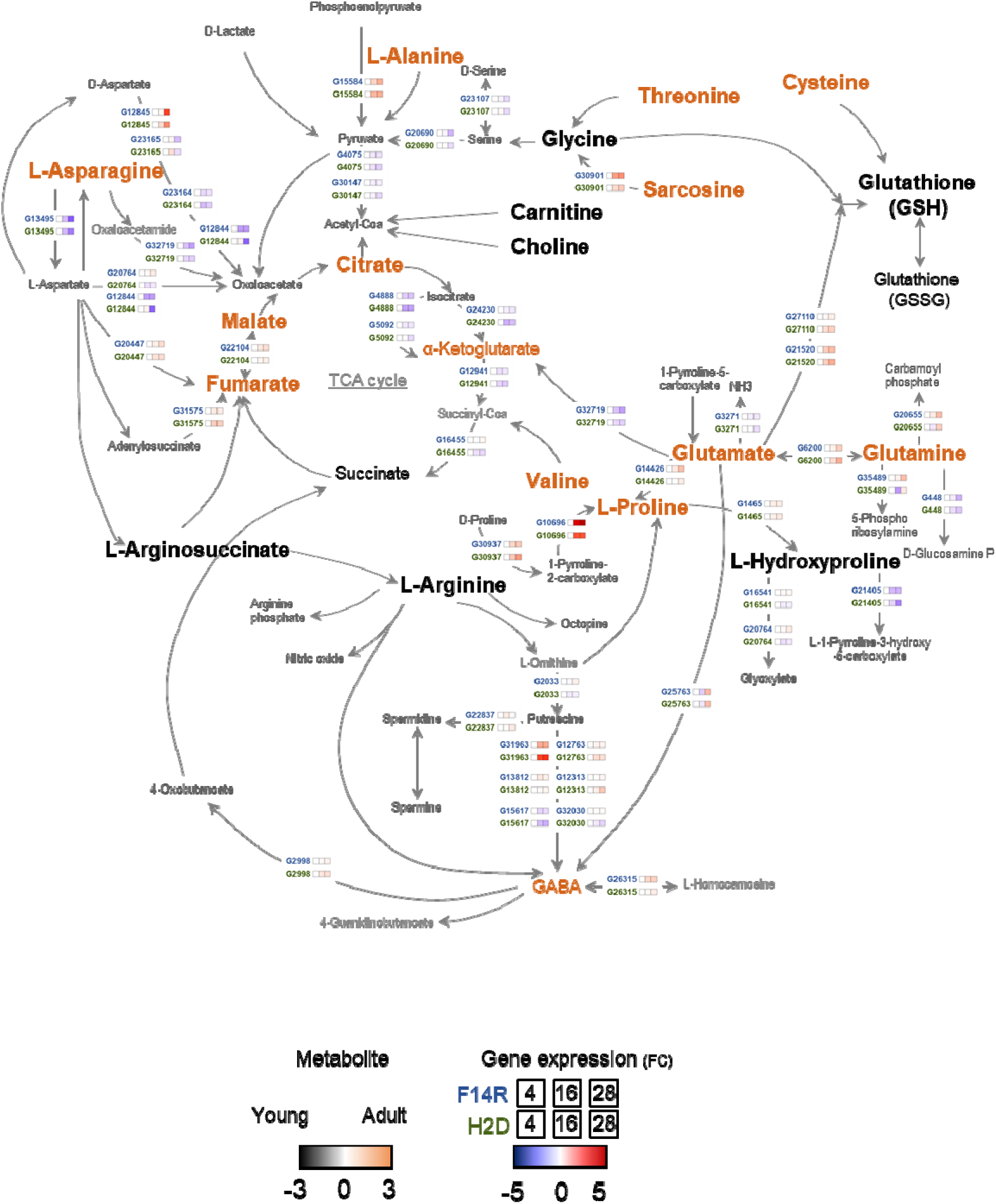
Metabolic map integrating transcriptomic and metabolomic data associated with age in F14R and H2D. Schematic representation of metabolite and gene expression changes associated with ageing. The map illustrates metabolites and corresponding gene expression changes linked to age. Metabolites that accumulate (higher abundance) in 28-month-old oysters are shown in orange, whereas those without accumulation (low abundance) are shown in black. Metabolites detected in both families are highlighted in bold. Genes are color-coded according to fold change (FC) at a false discovery rate (FDR) < 0.05, using 4-month-old oysters as the reference group F14R (Blue) and H2D (Green). Arrows represent direct metabolic conversions. Family-specific results are shown separately in Figure S7 (F14R) and Figure S8 (H2D).

The concordance between metabolite levels and corresponding gene expression patterns associated with younger and older conditions strongly supports our integrative multi-omics approach and underscores the biological relevance of the identified pathways in age-dependent metabolic modulation within the context of POMS in oysters (**Figures 6**, **S7,** and **S8**).

Integrating transcriptomic and metabolomic data revealed a coherent age-associated reprogramming of central carbon and amino acid metabolism, reflecting altered mitochondrial function during ageing in oysters. The accumulation of TCA cycle intermediates with age (metabolomics: citrate, malate, and fumarate) coincided with the down-expression of multiple downstream genes, such as the succinate-CoA ligase (ADP-forming) subunit β mitochondrial (G16455), the isocitrate dehydrogenase (NAD) subunit β mitochondrial (G24230), the isocitrate dehydrogenase (NADP) mitochondrial-like (G5092), and the isocitrate dehydrogenase (NADP) cytoplasmic (G4888). These enzymes are involved in mitochondrial oxidative phosphorylation, suggesting a slowdown of TCA cycle flux of energy and reduced metabolic turnover in older oysters. In parallel, increased glutamate and glutamine levels in older oysters suggested a shift toward nitrogen storage, likely reflecting reduced amino acid turnover due to diminished TCA cycle activity. This interpretation is supported by the under-expression of the mitochondrial glutamate dehydrogenase gene (G3271) identified in both families, a key link between amino acid metabolism and the TCA cycle. Interestingly, the accumulation of glutamate and glutamine did not lead to enhanced antioxidant capacity, as levels of both reduced (GSH) and oxidized (GSSG) glutathione were lower in older oysters compared to younger ones, indicating regulated antioxidant defense. In both families, we also observed accumulation of L-Proline, accompanied by reduced L-Hydroxyproline levels and strong over-expression of the trans-L-3-hydroxyproline dehydratase gene (G10696) in older oysters. This enzyme catalyzes the dehydration of hydroxyproline, a metabolite derived from proline hydroxylation, which may serve as an adaptive strategy to buffer oxidative stress and preserve cellular integrity under diminished metabolic capacity. Finally, we detected accumulation of L-Asparagine alongside marked under-expression of the isoaspartyl peptidase/L-asparaginase gene (G13495). This was associated with reduced levels of L-Arginosuccinate and L-Arginine, two essential intermediates in arginine biosynthesis. Notably, at least in the F14R, over-expression of the nitric oxide synthase gene (G23422), homologous to salivary gland NOS, suggests an increased demand for L-Arginine as a substrate for the nitric oxide (NO) production. NO is a key mediator of innate immune responses and redox signaling. The observed metabolic imbalance, characterized by excessive L-Asparagine retention, limited arginine biosynthesis, and regulation of NO-generating pathways, suggests a complex age-associated reprogramming of nitrogen metabolism.

Taken together, the integration of epigenomic, transcriptomic, and metabolomic datasets reveals a coherent, multi-layered regulatory program associated with ageing in oysters. Epigenetic modifications appear to act as upstream regulators that shape transcriptional landscapes, particularly in immune– and stress-related pathways, thereby influencing the organism’s ability to cope with OsHV-1 µVar. These transcriptional programs are further mirrored in metabolic reorganization, where shifts in central carbon, amino acid, and redox metabolism reflect the functional consequences of gene expression regulation. The convergence across omics layers demonstrates that age-dependent changes are tightly interconnected, together driving a progressive transition toward reduced viral and altered physiological state, influencing the permissiveness to POMS in oysters.

## Discussion

This study aims to elucidate the molecular mechanisms by which ageing in *Magallana gigas* influences the permissiveness to Pacific Oyster Mortality Syndrome (POMS). The older oysters consistently displayed improved survival and markedly reduced OsHV-1 μVar replication compared with younger individuals of the same genetic background. This age-dependent protective effect, observed across all four families, aligns with previous reports of reduced POMS-related mortality in adult oysters (Arzul et al., 2002; Peeler et al., 2012; Pernet et al., 2012; Dégremont, 2013). To investigate the molecular basis of age-dependent differences in permissiveness to POMS, we performed an integrative multi-omics analysis (epigenetics, transcriptomics, and metabolomics) on two genetically distinct oyster families with contrasted phenotypes regarding permissiveness to POMS (F14R and H2D).

### Epigenetic and transcriptomic remodeling drives age-related immune trade-offs, influencing POMS permissiveness in oysters

The role of epigenetic mechanisms in invertebrate’ innate immunity remains poorly understood, but growing evidence suggests that epigenetic regulation acts as a key modulator of immune responses in invertebrates, including oysters (Gómez-Díaz et al., 2012; Fallet et al., 2020; Gu et al., 2022). Our analysis revealed age-associated DNA methylation changes in genes involved in cellular processes, metabolism, environmental stress responses, and immune functions, consistent with a previous study highlighting the contribution of epigenetic modulation to POMS pathogenesis in oysters (Fallet et al., 2022). We found that ageing progressively reshapes the oyster DNA methylation landscape, with pronounced changes in immune-related genes. Notably, several Toll-like receptor (TLR) genes, key pattern recognition receptors, and the adaptor gene Myd88 all exhibited increased DNA methylation levels with age. This finding supports previous findings linking epigenetic modulation of the TLR–NF-κB–Myd88 signaling to resistant and susceptible phenotypes in the POMS context (Gawra et al., 2023; Valdivieso et al., 2025).

Moreover, genome-wide DNA methylation profiling has shown that the NF-κB pathway is epigenetically regulated during *Vibrio alginolyticus* infections in oysters (J. Li et al., 2024), supporting the idea that age-dependent methylation shapes innate immune signaling and contributes to the improved outcomes observed in adult oysters during POMS outbreaks (Tang et al., 2020). DNA methylation is already recognized as essential during early oyster development (Riviere et al., 2013), and its gradual remodeling during ageing is likely to be not only a marker of age but also a driver of transcriptional reprogramming (Saint-Carlier & Riviere, 2015). While direct correlations between methylation and gene expression in invertebrates are often weak or absent (X. Wang et al., 2014; Dixon & Matz, 2022), evidence suggests that DNA methylation interacts with other epigenetic mechanisms to regulate chromatin accessibility (Badeaux & Shi, 2013). Such interactions may profoundly shape the relationship between epigenetic and transcriptional regulation (Bogan et al., 2023), ultimately influencing oyster defense capacity against POMS, as supported by our findings. The lack of gene-to-gene biological correlations between DNA methylation and gene expression regulation is a recurrent observation in oysters and other invertebrates (Gavery & Roberts, 2010; Johnson et al., 2022). Previous studies have shown that different genetic backgrounds in oysters can strongly influence the expression of specific genes within multigenic families, leading to different gene-level signatures but ultimately converging on similar biological functions (de Lorgeril et al., 2020). Together, these findings support the view that integrative multi-omics approaches are more informative when interpreted at the functional pathway or process level, rather than focusing solely on one-to-one gene relationships.

Ageing was also accompanied by extensive transcriptional reprogramming, affecting key biological processes that influence permissiveness to POMS in oysters. Specifically, these transcriptomic changes included an enhanced DNA repair capacity (Zhao et al., 2023) and a reshaped immune landscape characterized by the over-expression of antiviral defense genes (e.g., Defense/Response to virus, Defense response to symbiont) and the under-expression of the NF-κB signaling components, including genes in the Toll signaling pathway, MyD88-dependent Toll-like receptor signaling, and cytokinesis-related processes. These processes have been widely reported in oyster immunity (Huang et al., 2019) and in POMS pathogenesis in oysters (Green & Speck, 2018; de Lorgeril et al., 2018, 2020; Duperret et al., 2025). The attenuation of gene expression involved in the NF-κB signaling in adults suggests a dampened inflammatory response (Rothschild et al., 2018), potentially reflecting a shift from broad pro-inflammatory programs in young oysters toward more targeted antiviral defenses in older oysters. In *M*. *gigas*, this pathway plays roles in developmental, cellular, and immune response processes (Yu et al., 2018), and it is mostly mediated by the TLR (L. Zhang et al., 2011). This immune reprogramming, together with the over-expression of virus recognition and response genes, may represent a strategy that enhances viral control while limiting inflammatory damage (de Lorgeril et al., 2020). Indeed, host immune gene expression dynamics are key determinants of infection outcome, influencing both pathogen proliferation and disease progression (Alizon et al., 2011). These dynamics are also observed from our results, where age-associated epigenetic and transcriptional remodeling shifts, enabling older oysters to exert stronger control over OsHV-1 μVar, whereas younger individuals remain highly susceptible, as reflected by higher mortality rates and viral replication. However, this ontogenetic shift in immune prioritization may involve a life-stage trade-off. Young oysters appear biased toward antibacterial defenses, while older ones may favor antiviral responses, as depicted with epidemiological observations that juveniles were more susceptible to OsHV-1 μVar, whereas adults were predominantly affected by *Vibrio aestuarianus* (Azéma et al., 2017; Green et al., 2016).

### Age-dependent redox and metabolic reprogramming shape immune responses and susceptibility to POMS

In bivalves, the immune defense mainly relies on phagocytosis by haemocytes by enzymatic or oxidative degradation (Canesi et al., 2002). Upon activation, the oyster immune cells generate reactive oxygen species (ROS), a highly reactive molecule produced as byproducts of mitochondrial respiration and enzymatic activity (Sussarellu et al., 2013). While ROS production is crucial for pathogen elimination, their excessive accumulation can damage host tissues (C. Li & Jackson, 2002), making their tight regulation essential. This balance is achieved through the action of antioxidant enzymes, which sustain the cellular redox state. In *M. gigas*, redox homeostasis has been identified as a central integrator of stress adaptation, linking antioxidant defenses with energy metabolism, immunity, and development (Trevisan & Mello, 2024). Our findings are consistent with this view, highlighting that redox regulation not only safeguards cellular integrity, but also shapes the metabolic and immune responses during oyster aging, therefore influencing permissiveness to POMS. Understanding the intricate relationship between immune, metabolic pathways, and redox balance is pivotal for environmental stress induced by pathogens in oysters, especially in the context of some redox-mediated immuno-metabolic adaptations that may alter the susceptibility.

The integration of transcriptomic and metabolomic data revealed age-associated regulation of glutathione metabolism, underscoring its central role in shaping the cellular redox environment and influencing susceptibility to POMS (Samain, 2011). Glutathione (GSH), a tripeptide composed of glutamate, cysteine, and glycine, is a key antioxidant that neutralizes reactive oxygen species (ROS) through reversible redox cycling between its reduced (GSH) and oxidized (GSSG) forms. An optimal GSH/GSSG ratio is critical for maintaining redox homeostasis, protecting macromolecules from oxidative damage, and supporting cellular detoxification (Trevisan & Mello, 2024). However, excessive reduction (reductive stress) can also be deleterious, disrupting redox-sensitive signaling and protein folding, and causing DNA damage (Xiao & Loscalzo, 2020). In our dataset, younger oysters exhibited higher concentrations of both GSH and GSSG, reflecting a more dynamic redox buffering system and greater antioxidant capacity at early life stages compared to older oysters. This redox flexibility may enhance protection against oxidative stress and support immune activation during OsHV-1 infection. However, at these early stages, the organisms remain largely unable to control the infection, resulting in higher mortalities. These findings are consistent with the observation that ageing is associated with the over-expression of genes involved in the DNA repair process, likely reflecting a compensatory mechanism to counteract cumulative oxidative damage and maintain genome integrity. Older oysters displayed lower GSH/GSSG levels, consistent with a less reducing intracellular environment and altered immune signaling, as reflected in their transcriptomic profiles in genes involved in response to virus. The metabolic shifts in glycine and glutamate in old oysters, which are necessary precursors of GSH, further indicate age-related remodeling of antioxidant defenses (Rebrin & Sohal, 2008). These adaptations likely fine-tune the nuclear factor kappa-light-chain-enhancer of activated B cells (NF-κB) signaling (Asehnoune et al., 2004; Deponte, 2013; Lingappan, 2018), a process also observed in the transcriptomic data, thereby enhancing immune responsiveness. The Nitric oxide (NO) metabolism also emerged as a redox-linked pathway modulated by age. NO is synthesized by nitric oxide synthases from L-arginine and serves as both a potent antimicrobial effector and a sentinel of environmental stress (Nakayama & Maruyama, 1998; Ottaviani et al., 1993; Rivero, 2006; Tafalla et al., 2003). In young oysters, we showed elevated L-arginine levels and higher nitric oxide synthase expression, suggesting enhanced NO production and immune responsiveness. Conversely, older oysters exhibited reduced L-arginine availability, accumulation of L-asparagine, and down-expression of isoaspartyl peptidase/L-asparaginase, an enzyme driving L-asparagine degradation into L-aspartate in the argininosuccinate–arginine pathway, indicating a reduced flux toward NO synthesis. Transcriptomic analyses also revealed that multiple biosynthetic and metabolic pathways, including lipid metabolism, catabolic processes, the TCA cycle, and ATP metabolism, were negatively associated with age. The metabolomic data further illustrated a rewiring of the TCA cycle in oysters, with fumarate, malate, citrate, and α-ketoglutarate accumulated in older oysters, whereas succinate was more abundant in younger ones. Malate, derived from aspartate, succinate or glucose, succinate pathways, enters the mitochondria and can be processed through two complementary routes: an oxidative branch, generating NADH and ATP via substrate-level phosphorylation and complex I–mediated proton pumping, or a reductive branch, which regenerates succinate and uses fumarate as an alternative electron acceptor under low oxygen conditions (Müller et al., 2012; Trevisan & Mello, 2024). These metabolic routes highlight the flexibility of oyster mitochondrial metabolism, which can shift between oxidative and reductive modes in response to changing energy demands. The observed metabolite patterns may suggest a reduced oxidative flux and energy-conserving metabolism in older oysters, contrasted with a more active aerobic profile in younger ones.

### mTOR signaling as a central hub linking metabolism, autophagy, and immune defense in oyster aging

Our integrated omics analyses point to the mechanistic target of rapamycin (mTOR) pathway as a central regulator of age-associated changes in the oyster physiology. The mTOR pathway is a well-established master regulator of growth, metabolism, and cellular senescence, integrating environmental and cellular signals to coordinate anabolic processes, mitochondrial activity, and stress responses (Weichhart, 2018; Papadopoli et al., 2019). The observed age-dependent decline in energy metabolism and altered redox regulation in old oysters may reflect changes in the mTOR signaling activity, as cellular energy and nutritional state directly regulate mTOR to control associated signaling pathways downstream (Q. Zhang et al., 2023) and autophagy in bivalves (Kim et al., 2011; Levine & Deretic, 2007). In the context of POMS, starved oysters enhanced autophagy, improving their capacity to cope with the OsHV-1 µVar infection by limiting viral replication (Pernet et al., 2019). In oysters, autophagy is functional and modulates OsHV-1 μVar proliferation (Leprêtre et al., 2021; Moreau et al., 2015; Picot et al., 2019, 2020, 2022). In this study, we identified several genes involved in the autophagy/mTOR pathway displaying age-related changes at the epigenetic and transcriptional level, such as the inhibitor of nuclear factor kappa-B kinase subunit alpha (IKKα, G1756) and rapamycin-insensitive companion of mTOR (Rictor, G270) in both families. In addition, we found 28 and 34 genes downregulated with age in F14R and H2D, respectively, that are implicated in the KEGG mTOR signaling pathway, underscoring widespread transcriptional repression of this central regulatory network with age. Together, these findings highlight that mTOR-associated signaling components are broadly modulated during oyster ageing, potentially altering metabolic regulation, immune signaling, and autophagy, and providing a molecular basis for age-dependent variation in permissiveness to POMS.

## Conclusion

This multi-omics study demonstrates that ageing in *Magallana gigas* is characterized by adaptive reprogramming that reduces permissiveness to Pacific Oyster Mortality Syndrome (POMS). By integrating epigenomic, transcriptomic, and metabolomic data, we show that epigenetic remodeling of key immune regulators, such as Toll-like receptors and MyD88, aligns with transcriptional rewiring of NF-κB and ubiquitin pathways, indicating fine-tuned control of innate immunity. Our data also provide strong evidence that mTOR signaling is repressed with age, likely promoting autophagy and enhancing viral control, thereby improving outcomes during OsHV-1 infection. This regulatory shift is tightly interconnected with metabolic reprogramming, including reduced tricarboxylic acid (TCA) cycle flux, remodeling of nitrogen metabolism, and altered glutathione dynamics, collectively supporting a stress-tolerant, energy-conserving phenotype. Together, these results reveal an evolutionary trade-off between growth and immune defense: juveniles’ metabolic activity increases their susceptibility to viral proliferation, whereas adults shift their resources toward cellular maintenance and antiviral preparedness, resulting in improved survival during POMS outbreaks.

## Data availability

RNA-seq data and EM-seq analyses have been made available through the ENA database under Bioproject DECICOMP: PRJEB86647.

EM-seq data were submitted under accession numbers PRJEB105019.

RNA-seq data were submitted under accession numbers PRJEB105132.

Metabolomic data have been made available through Metabolight database under the accession number MTBLS13463

Complementary information is available from the corresponding authors on reasonable request.

## Credit author statement

Funding acquisition: G.M.

Conceptualization: B.P., J.V.D, and G.M.

Methodology: B.P., J.V.D, and G.M

Investigation: A.V., L.D., E.T., A.L., J.V.D., and G.M.

Supervision: E.T., A.L., J.V.D., and G.M

Writing original draft: A.V., J.V.D.

Writing review and editing: all authors

## Acknowledgments

We would like to thank the staff of the Ifremer experimental platforms at Argenton and Bouin for providing experimental facilities and for the production of the biological material used in this study. We thank Bruno Chollet, Louis Boismorand, Charlotte Corporeau, Mathias Huber, Marianne Alunno-Bruscia and Moussa Diagne for assistance in the experimental design and procedure. We thank the Montpellier Alliance for Metabolomics and Metabolism Analysis (MAMMA) platform (BioCampus Montpellier) for technical support in the metabolomic analysis.

