## Supplementary figures for "Multi-Omics Reprogramming Drives a Counterintuitive Reversal of Disease Susceptibility During Ageing"

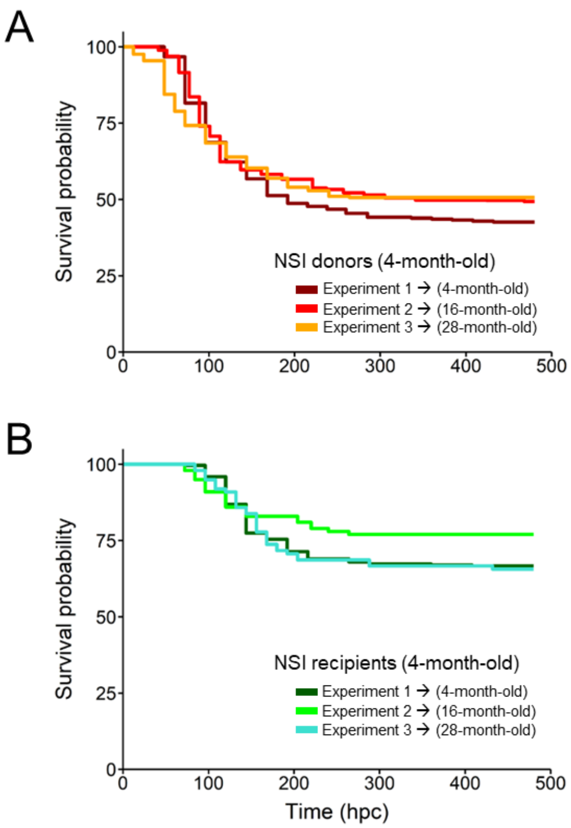


**Figure S1. Kaplan–Meier survival of donor oysters in sequential POMS challenges**

Kaplan–Meier survival curves showing **A)** 4-month-old donor NSI and **B)** 4-month-old NSI recipient oysters used in each POMS cohabitation challenge over three consecutive years: Experiment 1 (4-month-old, 2020), Experiment 2 (16-month-old, 2021), and Experiment 3 (28-month-old, 2022).

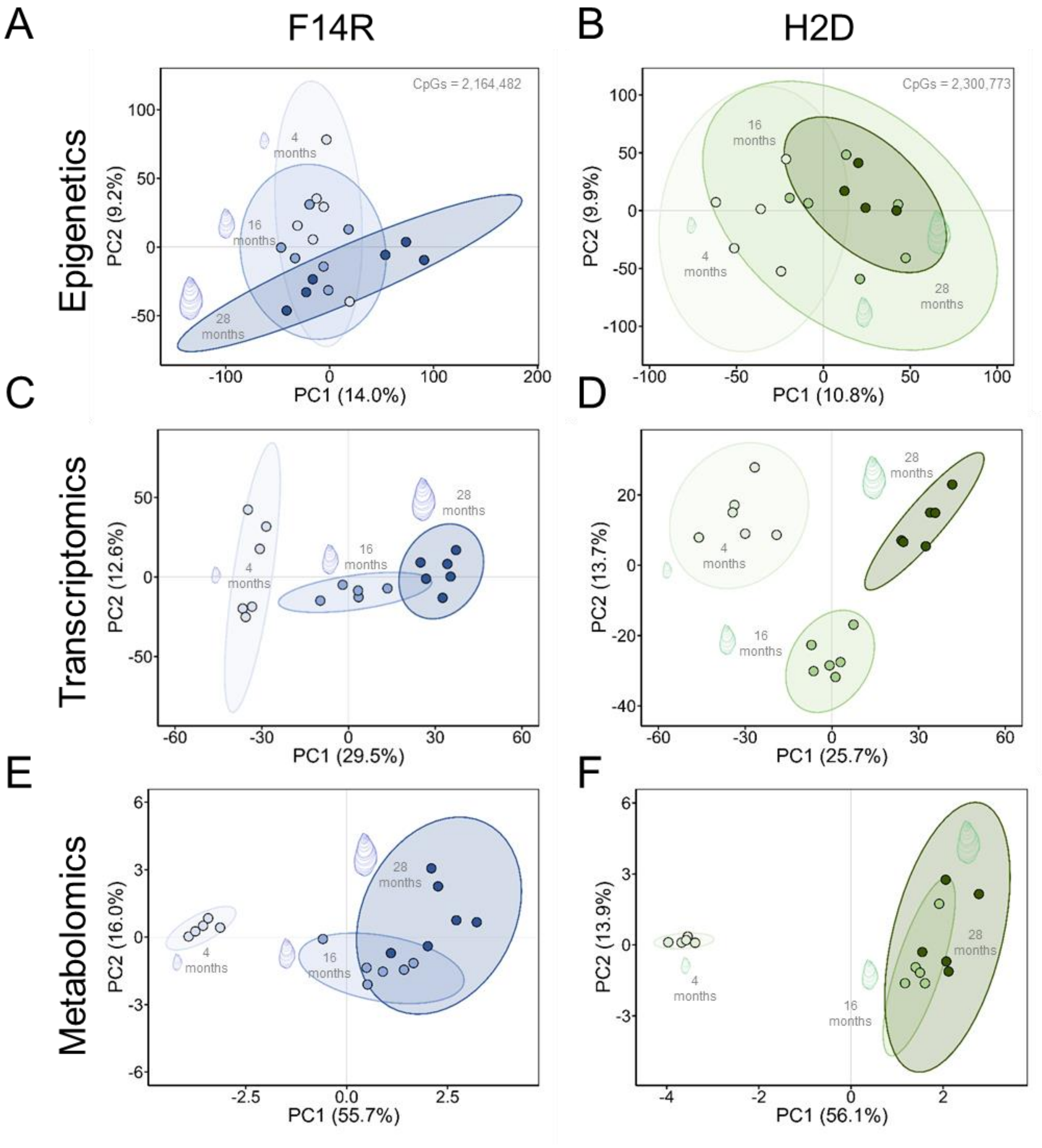


**Figure S2. Principal Component Analysis (PCA) in F14R and H2D families across 4-, 16-, and 28-month-old groups**

**A–B)** PCA of epigenetic (methylome) profiles based on informative CpG methylation (≥ 8 × coverage), including CpG sites common to 2,164,482 in F14R and 2,300,773 in H2D across age groups. **C–D)** PCA of transcriptomic (gene expression) profiles derived from normalized gene expression data (25,476 genes). **E–F)** PCA of metabolomic (metabolites) profiles based on normalized metabolite abundances (74 metabolites).

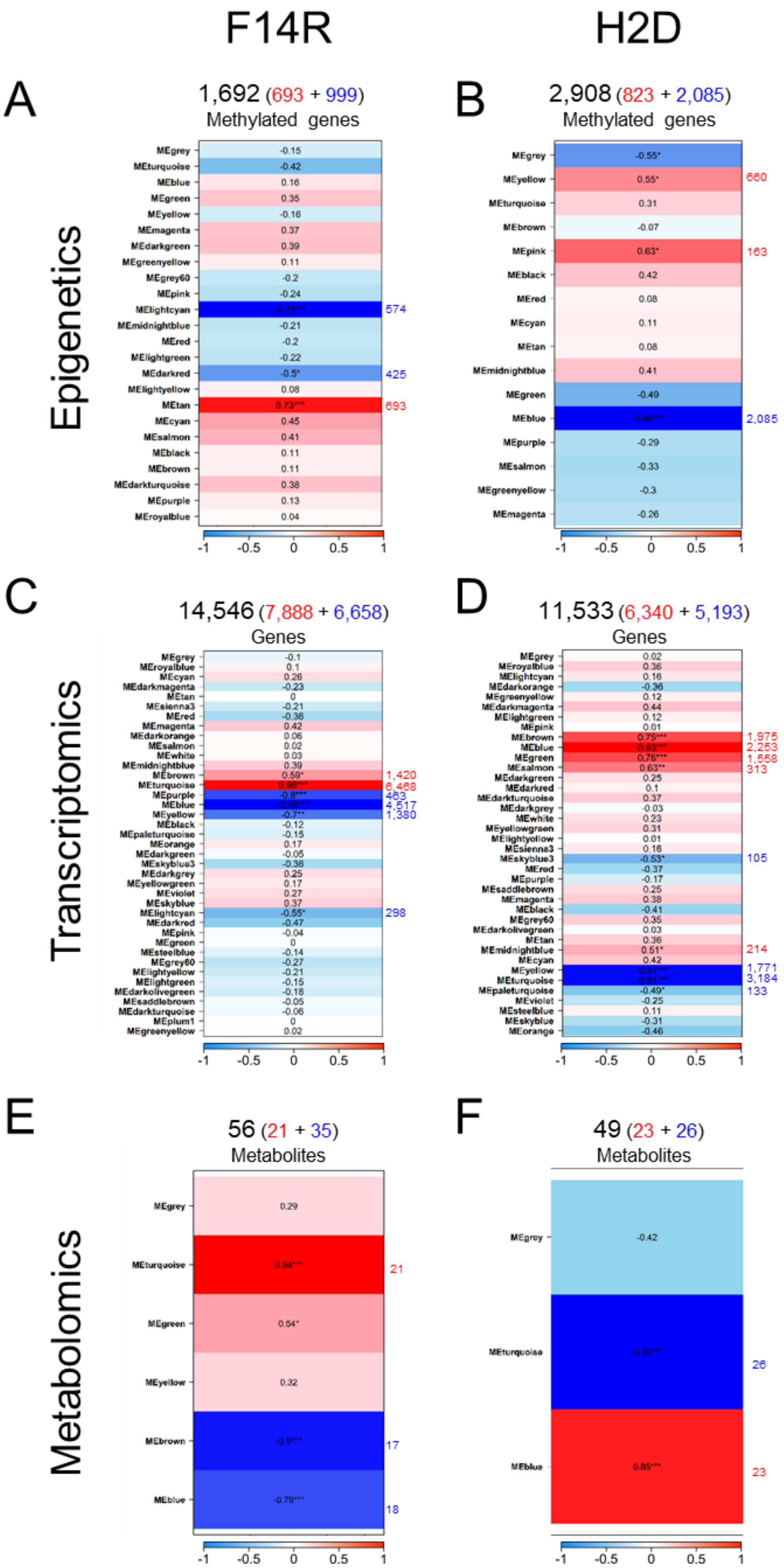
F14R
H2D
A
1,692 (693 + 999)
Methylated genes
B
2,908 (823 + 2,085)
Methylated genes
Epigenetics
C
14,546 (7,888 + 6,658)
Genes
D
11,533 (6,340 + 5,193)
Genes
Transcriptomics
E
56 (21 + 35)
Metabolites
F
49 (23 + 26)
Metabolites
Metabolomics

**Figure S3. Modules from WGCNA analysis showing the correlations with age in F14R and H2D families**

Modules showing positive (red; increased levels) and negative (blue; decreased levels) Pearson's correlations (r) with age in **A–B)** epigenetic (methylome), **C–D)** transcriptomic (gene expression), and **E–F)** metabolomic (metabolites) datasets. Only the significant correlations marked with asterisks in each module (* = P-value < 0.05; ** = P-value <0.01; and *** = P-value < 0.001) with the number of features within each module were considered for further analysis.

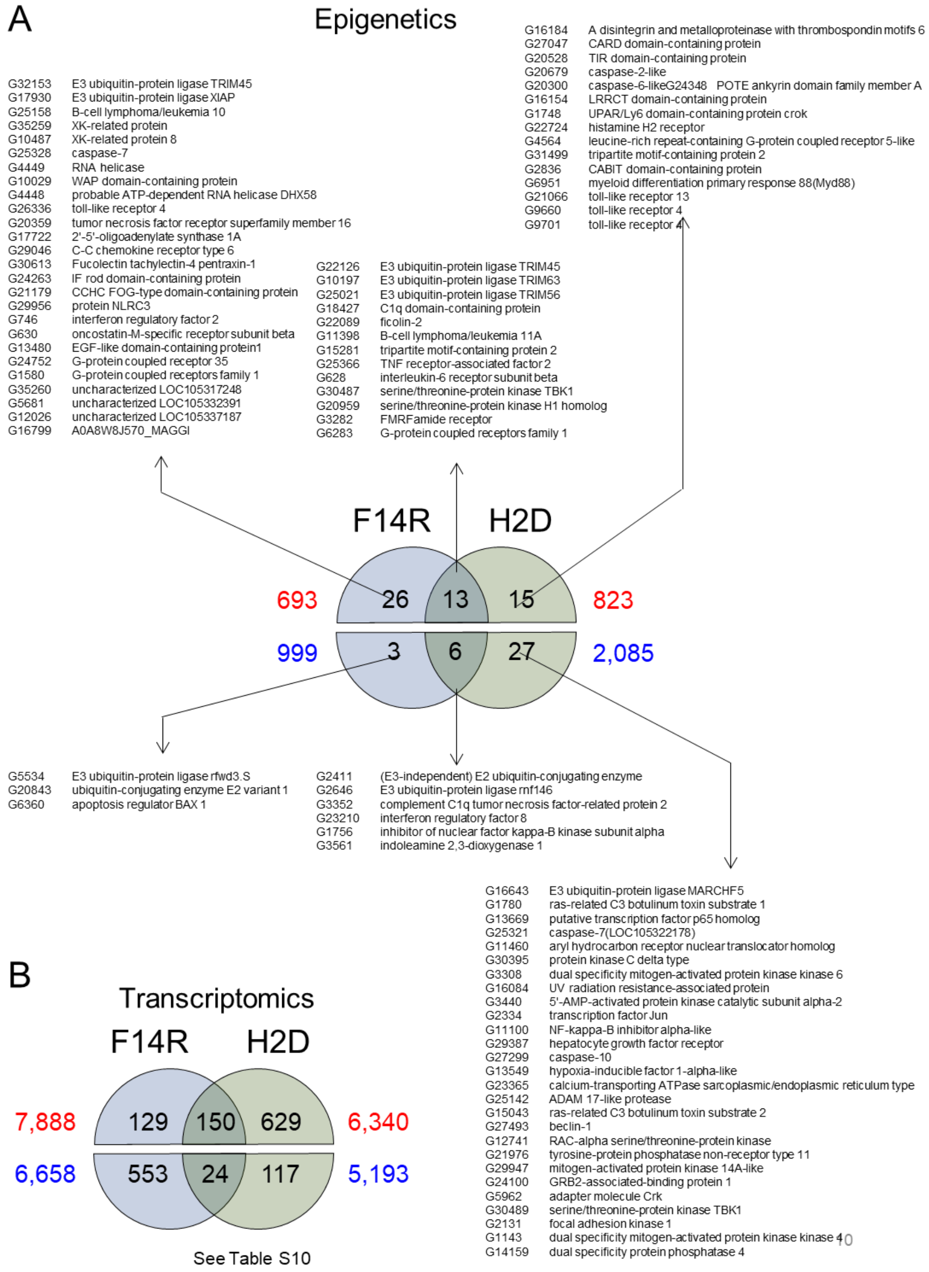


**Figure S4. Age-associated DNA methylation changes (Epigenetics) and gene expression (Transcriptomics) in genes involved in stimulus response and immune system processes.**

**A)** Common and specific genes between F14R and H2D with positive and negative DNA methylation changes associated with age involved in Gene Ontology (GO-terms) categories of Response to stimulus and Immune system process between F14R and H2D. **B)** Common and specific genes between F14R and H2D with positive and negative gene expression changes associated with age, involved in Gene Ontology (GO-terms) categories of Response to stimulus and Immune system process.

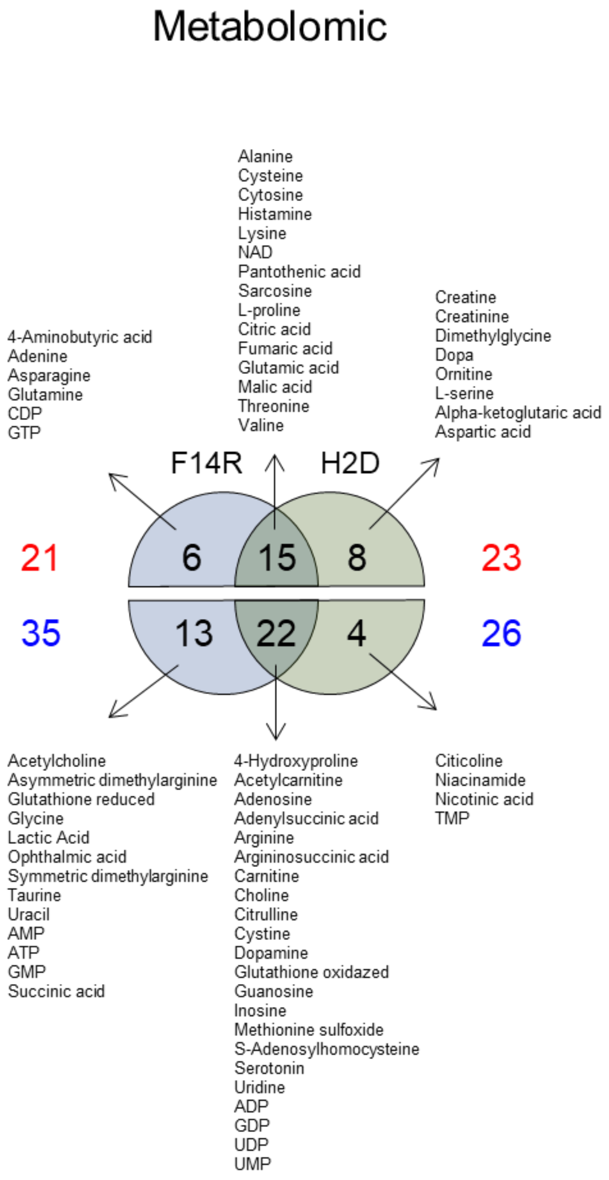


**Figure S5. Age-associated metabolites (Metabolomic)**

Common and specific metabolites between F14R and H2D with positive and negative changes associated with age.

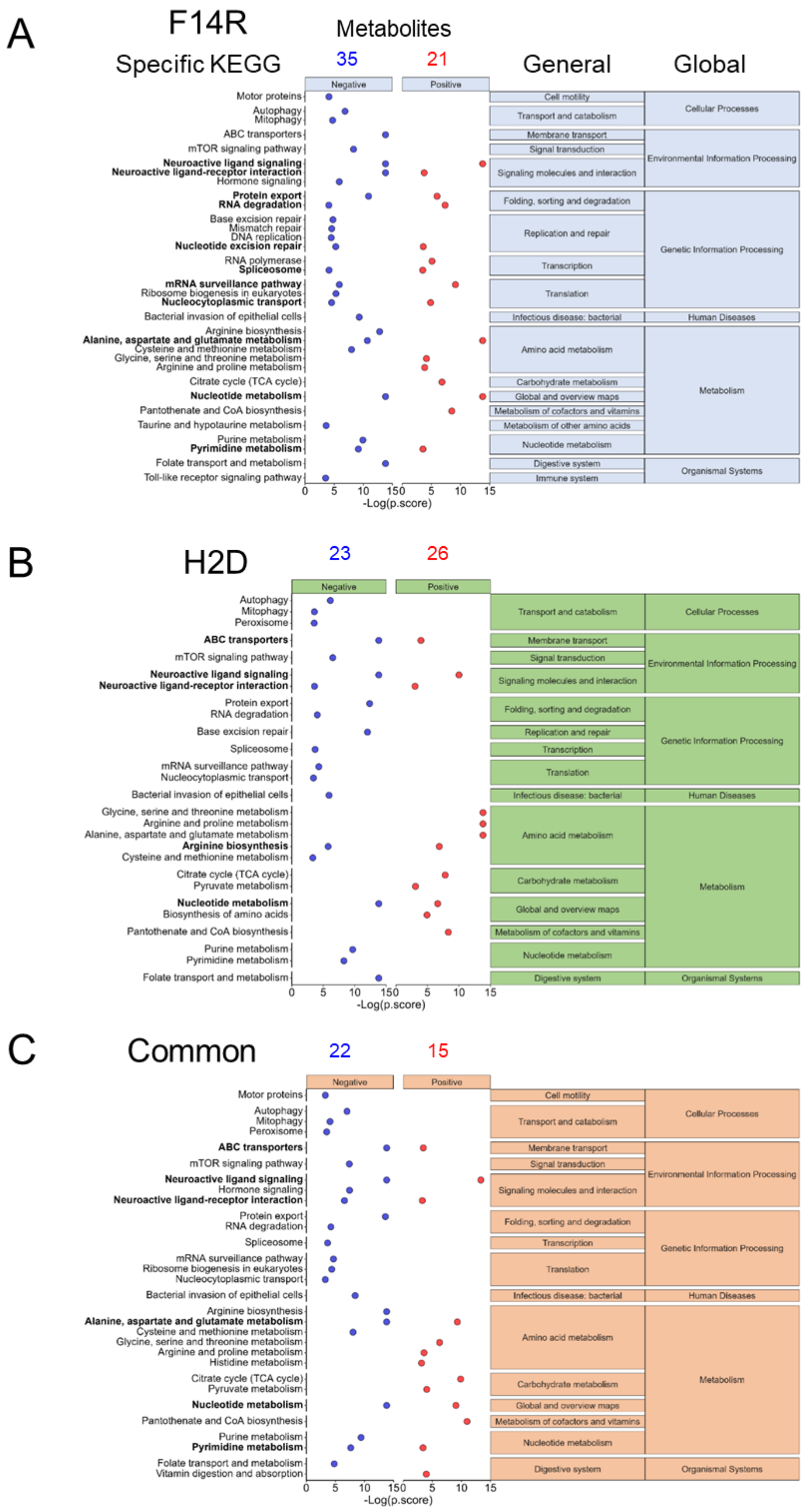
A
F14R
Specific KEGG
Metabolites
35
21
General
Global
Negative
Positive
Motor proteins
Autophagy
Mitophagy
ABC transporters
mTOR signaling pathway
Neuroactive ligand signaling
Neuroactive ligand-receptor interaction
Hormone signaling
Protein export
RNA degradation
Base excision repair
Mismatch repair
DNA replication
Nucleotide excision repair
RNA polymerase
Spliceosome
mRNA surveillance pathway
Ribosome biogenesis in eukaryotes
Nucleocytoplasmic transport
Bacterial invasion of epithelial cells
Arginine biosynthesis
Alanine, aspartate and glutamate metabolism
Cysteine and methionine metabolism
Glycine, serine and threonine metabolism
Arginine and proline metabolism
Citrate cycle (TCA cycle)
Nucleotide metabolism
Pantothenate and CoA biosynthesis
Taurine and hypotaurine metabolism
Purine metabolism
Pyrimidine metabolism
Folate transport and metabolism
Toll-like receptor signaling pathway
Cell motility
Transport and catabolism
Membrane transport
Signal transduction
Signaling molecules and interaction
Folding, sorting and degradation
Replication and repair
Transcription
Translation
Infectious disease: bacterial
Amino acid metabolism
Carbohydrate metabolism
Global and overview maps
Metabolism of cofactors and vitamins
Metabolism of other amino acids
Nucleotide metabolism
Digestive system
Immune system
Cellular Processes
Environmental Information Processing
Genetic Information Processing
Human Diseases
Metabolism
Organismal Systems
0 5 10 15 0 5 10 15
-Log(p.score)
B
H2D
23
26
Negative
Positive
Autophagy
Mitophagy
Peroxisome
ABC transporters
mTOR signaling pathway
Neuroactive ligand signaling
Neuroactive ligand-receptor interaction
Protein export
RNA degradation
Base excision repair
Spliceosome
mRNA surveillance pathway
Nucleocytoplasmic transport
Bacterial invasion of epithelial cells
Glycine, serine and threonine metabolism
Arginine and proline metabolism
Alanine, aspartate and glutamate metabolism
Arginine biosynthesis
Cysteine and methionine metabolism
Citrate cycle (TCA cycle)
Pyruvate metabolism
Nucleotide metabolism
Biosynthesis of amino acids
Pantothenate and CoA biosynthesis
Purine metabolism
Pyrimidine metabolism
Folate transport and metabolism
Transport and catabolism
Membrane transport
Signal transduction
Signaling molecules and interaction
Folding, sorting and degradation
Replication and repair
Transcription
Translation
Infectious disease: bacterial
Amino acid metabolism
Carbohydrate metabolism
Global and overview maps
Metabolism of cofactors and vitamins
Nucleotide metabolism
Digestive system
Cellular Processes
Environmental Information Processing
Genetic Information Processing
Human Diseases
Metabolism
Organismal Systems
0 5 10 15 0 5 10 15
-Log(p.score)
C
Common
22
15
Negative
Positive
Motor proteins
Autophagy
Mitophagy
Peroxisome
ABC transporters
mTOR signaling pathway
Neuroactive ligand signaling
Hormone signaling
Neuroactive ligand-receptor interaction
Protein export
RNA degradation
Spliceosome
mRNA surveillance pathway
Ribosome biogenesis in eukaryotes
Nucleocytoplasmic transport
Bacterial invasion of epithelial cells
Arginine biosynthesis
Alanine, aspartate and glutamate metabolism
Cysteine and methionine metabolism
Glycine, serine and threonine metabolism
Arginine and proline metabolism
Histidine metabolism
Citrate cycle (TCA cycle)
Pyruvate metabolism
Nucleotide metabolism
Pantothenate and CoA biosynthesis
Purine metabolism
Pyrimidine metabolism
Folate transport and metabolism
Vitamin digestion and absorption
Cell motility
Transport and catabolism
Membrane transport
Signal transduction
Signaling molecules and interaction
Folding, sorting and degradation
Transcription
Translation
Infectious disease: bacterial
Amino acid metabolism
Carbohydrate metabolism
Global and overview maps
Metabolism of cofactors and vitamins
Nucleotide metabolism
Digestive system
Cellular Processes
Environmental Information Processing
Genetic Information Processing
Human Diseases
Metabolism
Organismal Systems
0 5 10 15 0 5 10 15
-Log(p.score)

**Figure S6. KEGG pathway enrichment of age-associated metabolites.**

KEGG pathways from age-associated metabolites KEGG terms are displayed on the left, while broader functional categories (General and Global) are shown on the right. Enrichment is represented as −log(p-score) with negative (blue) and positive (red) age associations in **A)** Family F14R, **B)** Family H2D, and **C)** metabolites common to both F14R and H2D families.

# F14R

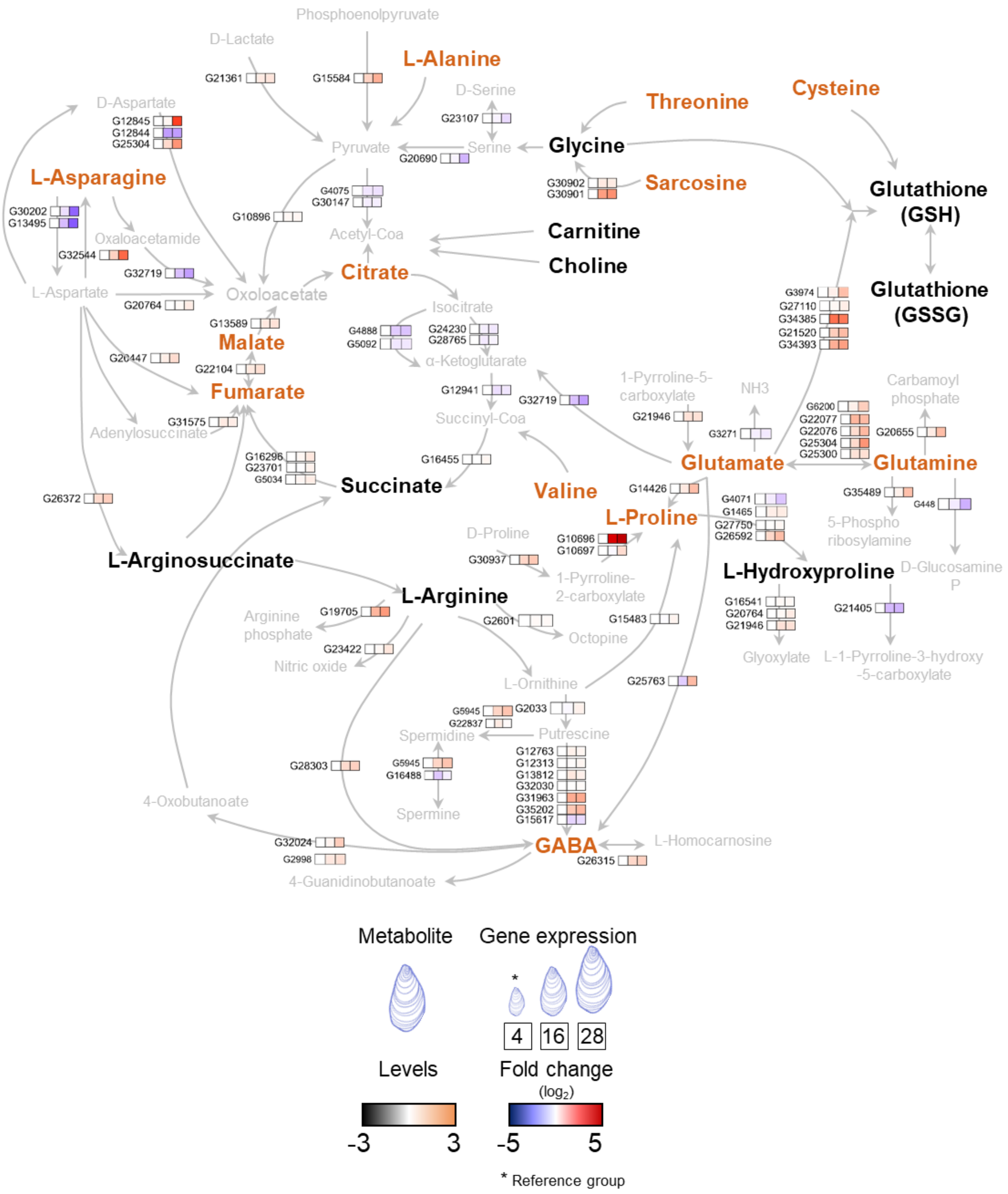


**Figure S7. Metabolic map integrating transcriptomic and metabolomic data associated with age in F14R.**

Schematic representation of metabolite and gene expression changes associated with ageing in F14R. The map illustrates metabolites and corresponding gene expression changes linked to age. Metabolites that accumulate (higher abundance) in 28-month-old oysters are shown in orange, whereas those without accumulation (low abundance) are shown in black. Genes are shown according to fold change (FC) at a false discovery rate (FDR) < 0.05, using 4-month-old oysters as the reference group F14R. Arrows represent direct metabolic conversions.

# H2D

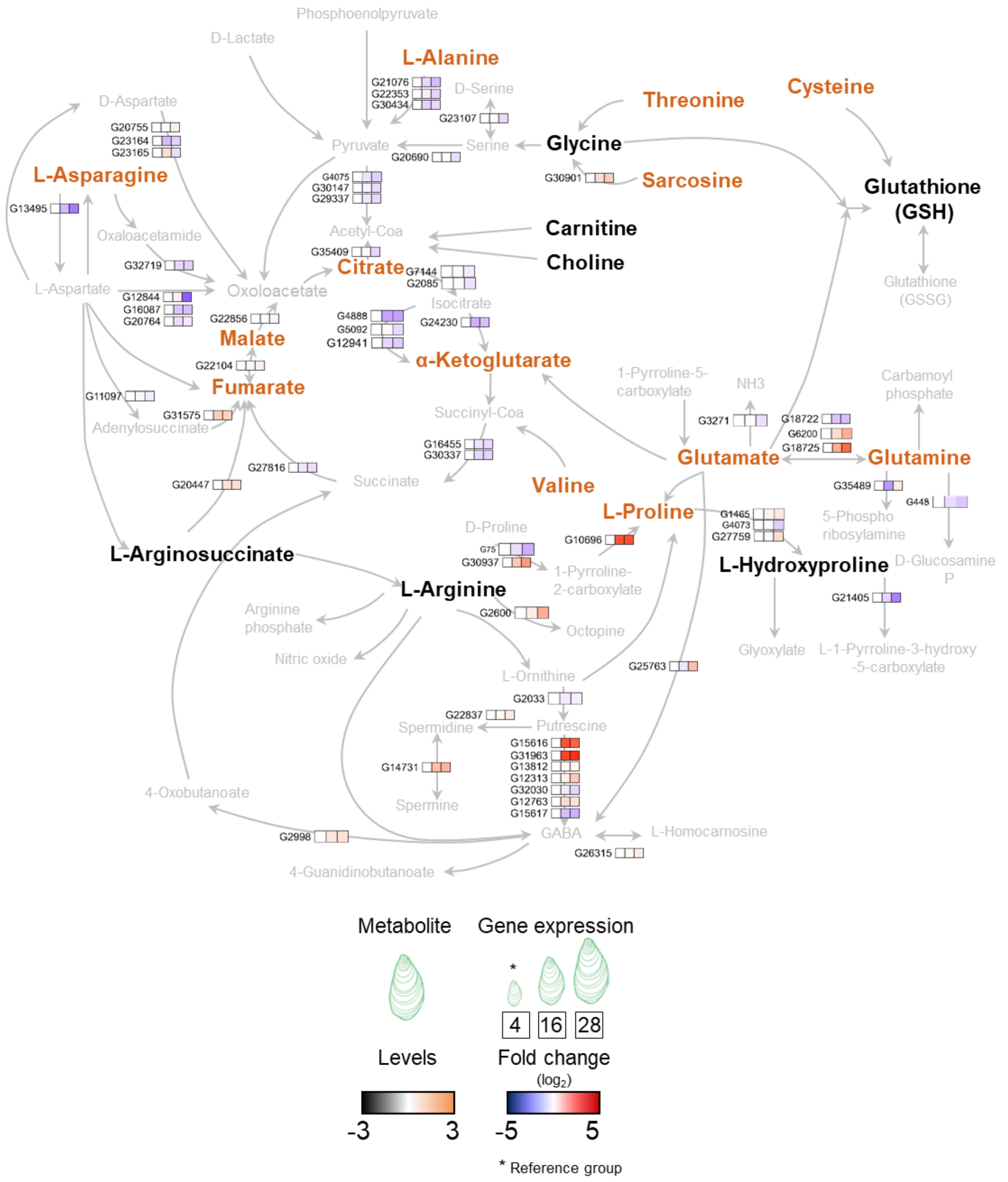


**Figure S8. Metabolic map integrating transcriptomic and metabolomic data associated with age in H2D.**

Schematic representation of metabolite and gene expression changes associated with ageing in H2D. The map illustrates metabolites and corresponding gene expression changes linked to age. Metabolites that accumulate (higher abundance) in 28-month-old oysters are shown in orange, whereas those without accumulation (low abundance) are shown in black. Genes are shown according to fold change (FC) at a false discovery rate (FDR) < 0.05, using 4-month-old oysters as the reference group F14R. Arrows represent direct metabolic conversions.
